# An improved whole life cycle culture protocol for the hydrozoan genetic model *Clytia hemisphaerica*

**DOI:** 10.1101/852632

**Authors:** Marion Lechable, Alexandre Jan, Brandon Weissbourd, Julie Uveira, Loann Gissat, Sophie Collet, Laurent Gilletta, Sandra Chevalier, Lucas Leclère, Sophie Peron, Carine Barreau, Régis Lasbleiz, Evelyn Houliston, Tsuyoshi Momose

## Abstract

The jellyfish species *Clytia hemisphaerica* (Cnidaria, Hydrozoa) has emerged as a new experimental model animal in the last decade. Favorable characters include a fully transparent body suitable for microscopy, daily gamete production and a relatively short life cycle. Furthermore, whole genome sequence assembly and efficient gene editing techniques using CRISPR/Cas9 have opened new possibilities for genetic studies. The quasi-immortal vegetatively-growing polyp colony stage provides a practical means to maintain mutant strains. In the context of developing *Clytia* as a genetic model, we report here an improved whole life cycle culture method including an aquarium tank system designed for culture of the tiny jellyfish form. We have compared different feeding regimes using *Artemia* larvae as the food and demonstrate that the stage-dependent feeding control is the key for rapid and reliable medusa and polyp rearing. Metamorphosis of the planula larvae into a polyp colony can be efficiently induced using a new synthetic peptide. The optimized procedures detailed here make it practical to generate new genetically modified *Clytia* strains and to safely maintain their whole life cycle in the laboratory.

## Introduction

Choice of model species is a fundamental decision in biological research. Standard models such as *Drosophila*, mice, or *Caenorhabditis elegans* benefit from clear advantages of accumulated knowledge, established laboratory strains, and experimental techniques. Non-standard model animals can also contribute, for instance, by providing insights into animal evolution, diversity and their interactions with the natural environment, as well as access to some biological processes absent or difficult to study in standard models (Cook et al., 2016; Goldstein and King, 2016). When using non-standard models, one significant hurdle is to establish standardized methods for raising and maintaining animals in the laboratory and controlling reproduction to obtain reliable embryonic and post-embryonic developmental stages. We describe here a robust laboratory culture method for the hydrozoan jellyfish *Clytia hemisphaerica* (Cnidaria), which has now become an accessible animal model to address a wide range of biological questions (Houliston et al., 2010; Leclère et al., 2016).

Cnidaria diverged early during animal evolution from the clade Bilateria, which includes almost all standard experimental model species. Adult bodies of most cnidarians are organized radially and comprise only two (diploblast) rather than three (triploblast) germ layers. They nevertheless possess nervous systems, muscles, and sensory organs that have evolved in parallel to those in found in Bilateria. Comparative studies between cnidarians and bilaterians are thus providing insights into how metazoans have acquired their current diversity. A variety of cnidarian species has been employed for experimental studies, in particular for developmental biology and evo-devo research. Notable examples are the anthozoan *Nematostella vectensis* (sea anemone) and the hydrozoan polyp species *Hydra (*freshwater) and *Hydractinia* (saltwater), (Galliot, 2012; Plickert et al., 2012). Laboratory culture of these three species is simplified by the absence of a pelagic medusa (jellyfish) stage, a life cycle stage characteristic of many cnidarian species in Medusozoa (including Hydrozoa, Scyphozoa, Cubozoa and Staurozoa) but absent in its sister clade Anthozoa. Indeed, culturing the medusa stage in the laboratory is challenging. Scyphozoan jellyfish of the popular species *Aurelia aurita* (Gold et al., 2019) are easy to generate from polyps and to maintain in aquaria, but the life cycle is long and difficult to complete (Spangenberg, 1965). A full life cycle has also been obtained in captivity for the direct developing (ie polyp-free) scyphozoan jellyfish, *Pelagia noctiluca* (Lilley et al., 2014; Ramondenc et al., 2019). In contrast, hydrozoan medusae are generally smaller with shorter lifespans, and thus attractive as laboratory jellyfish models.

We have developed *Clytia hemisphaerica* (Linnaeous, 1767) as a jellyfish model, primarily motivated by its suitability for developmental biology and cell biology. *Clytia* is transparent throughout the whole life cycle (Fig.1), an advantageous feature for microscopic observation. Daily release of gametes from jellyfish (Fig.1A) can be controlled simply by the dark-light cycle. The life cycle can be completed in as little as 2 months in the laboratory. Fertilized eggs develop to form a mature planula larva in 3 days (Fig.1B). Upon appropriate bacterial cues, conveniently substituted by adding synthetic neuropeptides GLWamide in sea water, they settle on a suitable substrate and metamorphose into primary polyps (Fig.1C). From each primary polyp, a colony composed of polyps specialized for feeding (gastrozooids) and budding medusae (gonozooids) then develops vegetatively by extension of the connecting stolon network (Fig.1D). Juvenile medusae (Fig.1E) bud continuously from the gonozooids if the colony is well fed, and reach adult size (about 10 mm bell diameter) and sexual maturity in 2-3 weeks in aquarium conditions (Fig.1A).

**Figure 1.**
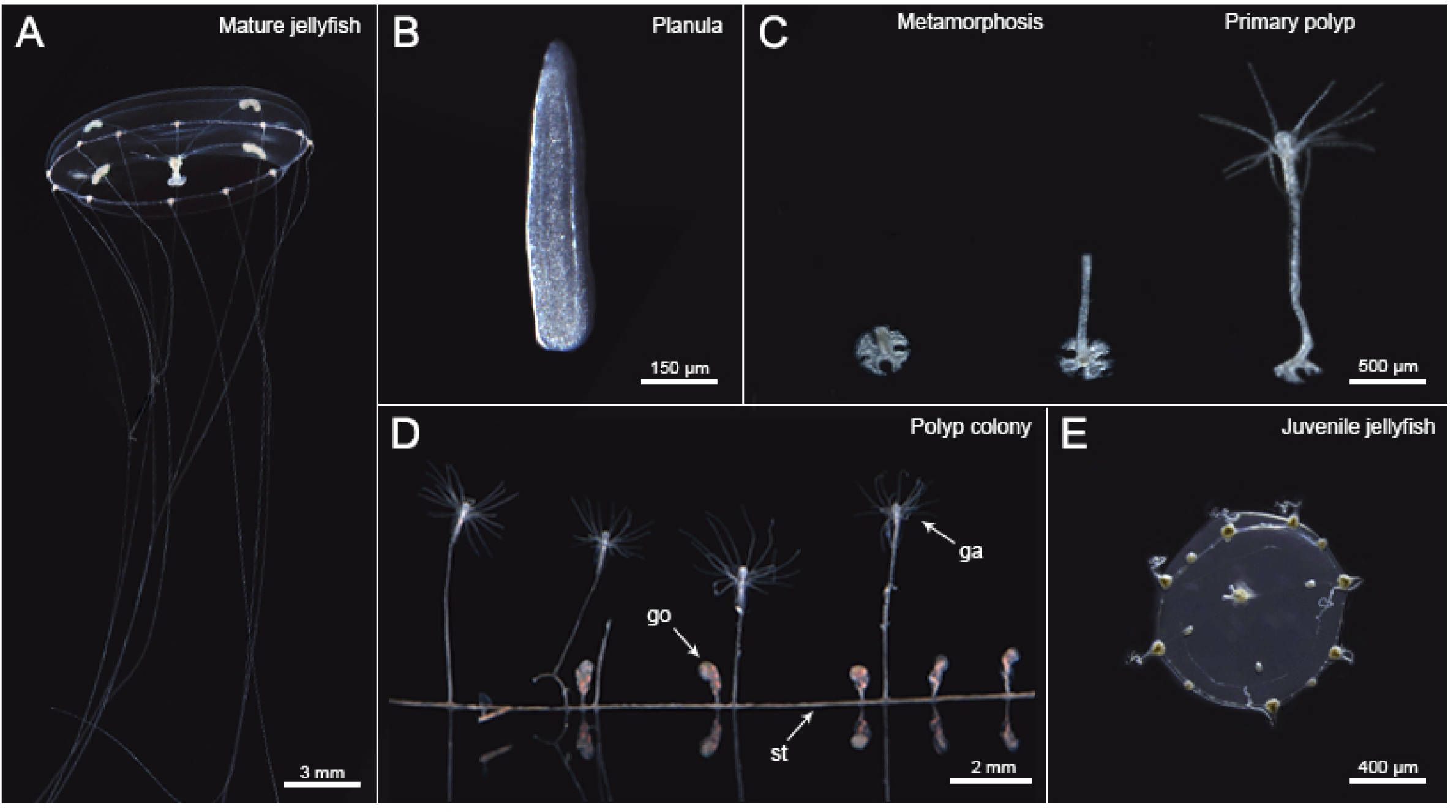
Stages of the *Clytia hemisphaerica* life cycle. *Clytia* life cycle consists of polyp stage, jellyfish stage and embryo-planula larva stage. (A) Adult jellyfish survives several weeks in the laboratory and spawns eggs or sperm, upon light illumination. (B) Fertilized eggs develop into planula larvae. (C) Planula undergoes metamorphosis when it recognizes a solid surface covered by biofilm, which can be induced by GLW-amide neuropeptide added to sea water in the laboratory conditions. (D) A primary polyp extends a stolon (st) horizontally and make a colony with multiple feeding polyps (ga:gastrozooids) and medusa-budding polyps (go: gonozooid) with sufficient feeding, which are all genetically clonal. € Gonozooid forms medusae buds, which are detached as juvenile jellyfish.

Available molecular and genetic resources include a whole genome sequence assembly and staged transcriptomes (Leclère et al., 2019; available on Marimba database http://marimba.obs-vlfr.fr). Efficient methods for gene function analysis have made *Clytia* an attractive genetic animal model, notably Morpholino antisense oligo and mRNA micro-injection into eggs prior to fertilization (Momose and Houliston, 2007), and more recently highly efficient CRISPR/Cas9-mediated gene KO (Momose and Concordet, 2016; Momose et al., 2018).

Here we detail the laboratory culture conditions and parameters affecting growth and development of *Clytia* over the whole life cycle. The culture system and tanks have been optimized for each stage. Daily feeding of *Artemia salina* nauplii is sufficient to maintain polyp colonies and medusae. We show how appropriate feeding regimes are critical factors for rapid growth in the laboratory, especially at very early medusa and primary polyp stages. The reliable culture system described here provides the essential basis for genetic studies in *Clytia*, for instance using mutants generated by CRISPR/Cas9 or emerging transgenesis technologies.

## Materials and methods

### Overview and rationale of the aquarium system for *Clytia* culture

Detailed *Clytia* life cycle culture material and methods are provided in the supplementary protocol; here we outline the culture concept. Tank design was found to be critical for stable *Clytia* culture, and we optimized different culture tanks for each life cycle stage (Fig. 2). Continuous water flow must be ensured to maintain the planktonic *Clytia* medusae (Fig.1A) in the water column. They remain in the bottom and die without water flow. We originally used 5-liter beakers with horizontal water rotation created by a paddle and a geared motor (5∼6 rpm), which was easy to set up but required intensive maintenance if used for large scale culture (see supplementary protocol). We thus devised a simple, closed-circuit aquarium system (Fig.3) that accommodates both polyp colonies and medusae.

**Figure 2.**
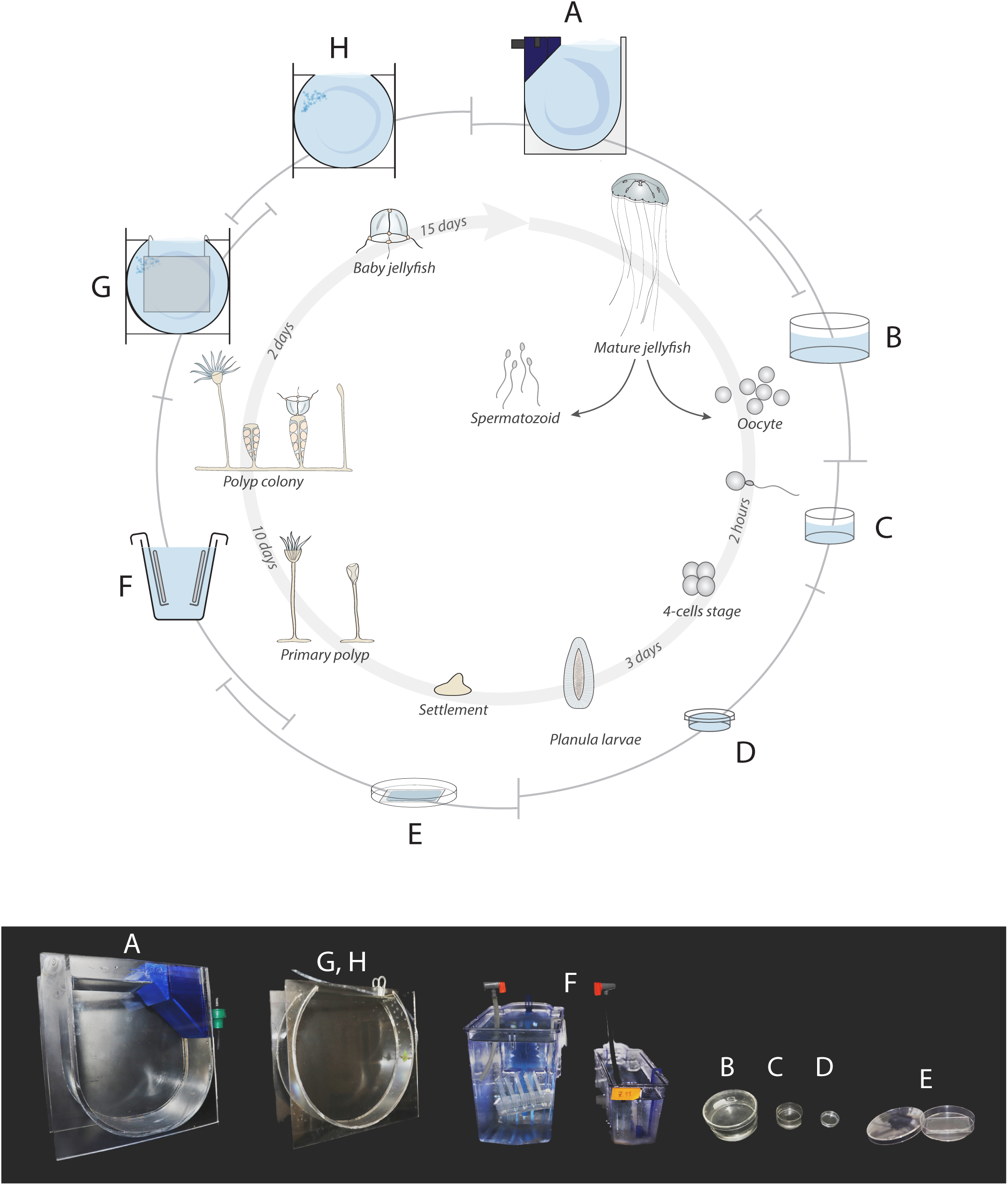
Use of different culture tanks optimized for different stages of the *Clytia* life cycle. (A) Mature jellyfish are maintained in Kreisel tanks (250×250×100mm). (B) Male and female medusae can be transferred into crystallizing dish (100 mm in diameter, 50 mm in depth) 1 to 2h before spawning and maintained on a shaker (50∼70 rpm). (C) Collected oocytes and sperm are transferred into smaller dishes (50 or 30 mm in diameter) to achieve fertilization. (D) Two to three hours later, developing embryos are transferred into small agarose-lined petri dishes. (E) Metamorphosis of planula larvae (60 hours after fertilization or older) is induced on a glass slide (75×50 mm) placed in a plastic petri dish (100 mm in diameter). (F) Glass slides are transferred into polyp tanks once primary polyps have completed development (standard zebrafish tank 280×150×100 mm; 3.5 liter or 280x 100x 60 mm;1.1 liter) and maintained to grow polyp colonies. (G) Juvenile medusae budded from gonozooid polyps are collected by leaving glass slides in nursery tank (280 mm diameter × 90 mm) or in crystallizing dish for up to 2 days. (H) The juvenile medusae are maintained in nursery tank or in crystallizing dish until their bell diameter reaches to 2.5 mm, large enough transfer to the Kreisel tank. See supplementary protocol for the detailed culture method.

**Figure 3.**
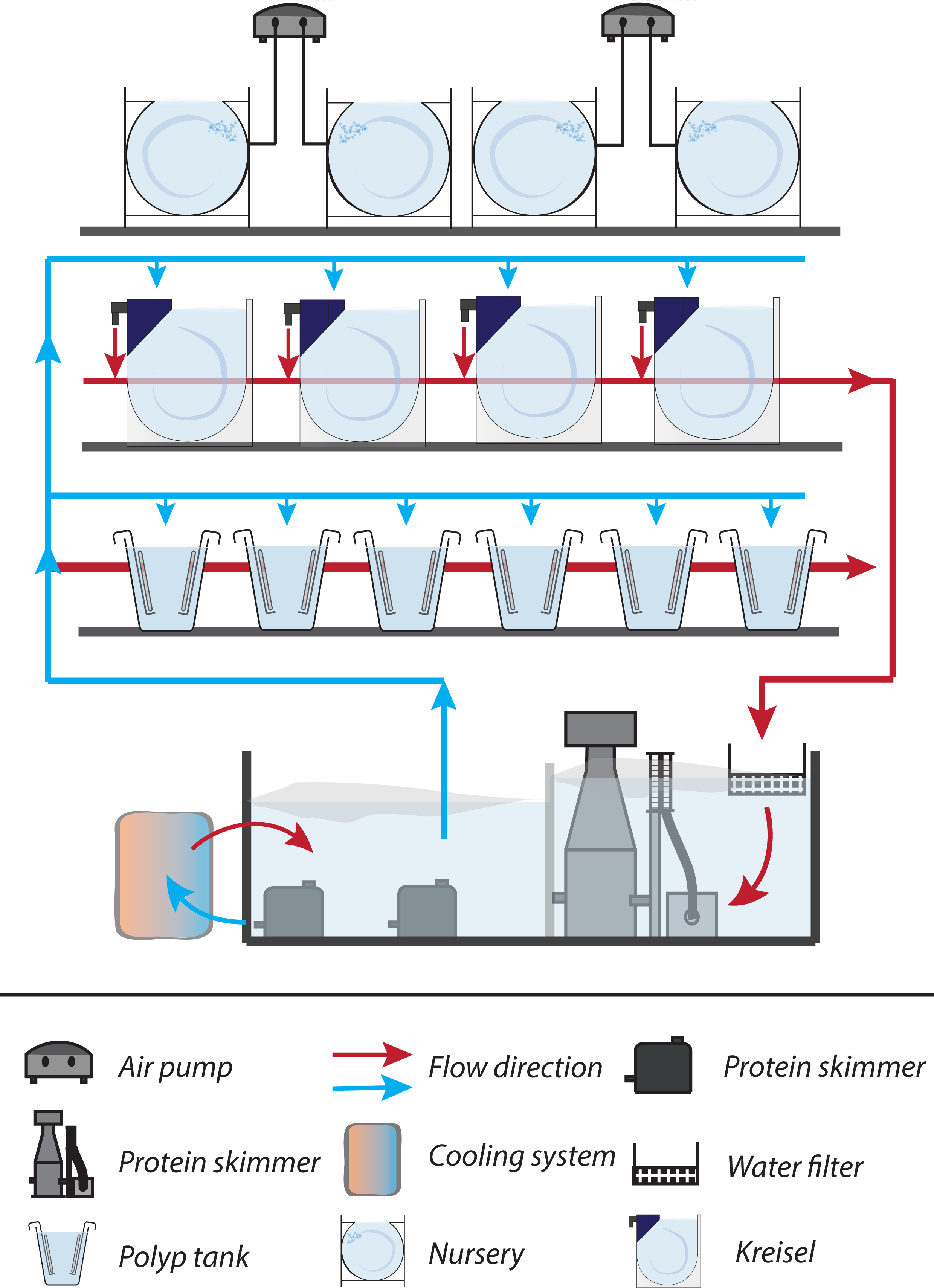
Culture system in the laboratory. Both Kreisel tanks (large medusae) and polyp tanks are integrated in a closed-circuit water aquaculture system. Water is supplied to each tank from the reservoir (50∼80 liters) by a submerged pump. Excess artemia, food leftovers or released *Clytia* jellyfish are primarily removed by a nylon mesh (200 µm) or filter pad (50 µm), then by protein skimmer, before the water returns to the reservoir. The temperature is adjusted by water chiller/heater. The salinity is measured every 2∼3 days and adjusted by adding deionized water. Roughly half of the water in the reservoir tank is replaced every 2 months.

In order to generate embryos, medusae can be temporary stored in 10 cm crystallizing dishes (Fig.2B) on a rotary shaker to collect eggs and sperm. For fertilization, collected gametes are mixed in smaller dish (3.5 cm diameter Fig.2C) and embryos are cultured in a 2% agarose-coated plastic dish (1.5∼3.5 cm diameter Fig.2D) until reaching the planula larva stage (Fig.1B). Planula larvae will undergo metamorphosis (Fig.1C) on glass slides (Fig.2E) and form polyp colonies (Fig.1D), which are maintained in polyp tanks (Fig.2F). Juvenile medusae are formed by budding from these colonies (from gastrozooid polyps) and are constantly released into the sea water. They are collected by keeping colony plates in a drum shaped “nursery tank” (Fig.2G), in which water rotation is created by air bubbles, or in dishes (Fig.2B). They are then grown to 2.5 mm bell diameter size in the same tank after removing polyp colonies (Fig.2H), at which point they are large enough to transfer into the “Kreisel tank” (Fig.2H). Kreisel tank (Fig.2A) is a key aquarium element of jellyfish culture system (Greve, 1968; Raskoff et al., 2016). We reduced the cost of the Kreisel tanks by combining a simple U-bottom tank in poly(methyl methacrylate) with 3D printed filter and nozzle parts (PET-G Ultra, Volumic 3D).

For both polyp and medusa stages, *Artemia* nauplii (1 day or older after hatching) are used for feeding (Fig.S3). We used *Artemia salina* (Sep-Art Ocean Nutrition) and smaller *Artemia franciscana* (Vinh Chau pond strain, Vietnam). Primary polyps and very young colonies are fed with smashed *Artemia*, until a stable colony of multiple gastrozooids has been established. After this colonies are fed regularly with live *Artemia* nauplii (once or twice a day, with minimum 6 hours of interval).

### Animal strains

Currently available wildtype strains are listed in Table 2. The main Z-series strains, originated from a single hermaphroditic Z strain, have been used in the *Clytia* user communities, including Z4B (female) and Z4C2 (male). Z4C2 strain male was used for whole genome sequencing (Leclère et al., 2019).

### Measurement of medusa growth

Juvenile Z4B strain medusae newly released from the colony were collected in the nursery tank (up to 1000 medusae in a 4-liter tank in one night, depending on the strain and colony size). After removing the polyp colony glass plate, juvenile medusae were kept in the tank until they reached about 2.5 mm in bell diameter, before being transferred to a Kreisel tank. The density of medusae was reduced to 50∼60 when they reached 5 mm in diameter. The bell diameter of 10 randomly sampled jellyfish was measured using an Axiozoom microscope using Zen software (Zeiss). For direct comparison of the effect of food types on early medusa growth, the same number of newly released medusae (115 medusae/tank) were placed in two nursery tanks and cultured for 6 days, feeding twice a day with about 350 artemia/tank (corresponding to 3 artemia/medusa). The bell size of all medusae in the tank was measured on the sixth day.

### Measurement of colony growth

Transplants of the Z4B strain colony grown on standard glass slides were cleaned so that only a single stolon-connected colony with 5 gastrozooids remained on each slide. The colonies were fed once or twice a day. The colony size was quantified every second day for two weeks as the number of functional feeding gastrozooids and gonozooids.

### Testing the efficiency of metamorphosis and early colony propagation

Metamorphosis of planula larvae was induced by transferring them onto glass slides covered with 0.1 ml/cm^2^ of seawater containing synthetic neuropeptides (Fig.2E). A selection of synthetic amidated peptides derived from the neuropeptide precursors ChePP2 and ChePP11 (Takeda et al., 2018) were tested as metamorphosis inducers (Table 2) on 3-day-old planula larvae obtained by crossing Z13 (male) and Z11 (female) strains. Planula attach firmly onto the slide within a few hours (Fig.1C left). Metamorphosis efficiency was measured by successful settlement on the slide at 12 hours, as the timing of stalk extension is variable, from several hours to days. To measure the survival and colony-formation rate of primary polyps, 40 successfully settled planulae were selected for each condition and the following development was monitored. The state of colony formation was classified into four categories: (1) initial state up to stalk formation (Fig.1C center), (2) successfully formed primarily polyp with tentacles (Fig.1C right), (3) colony stage with extended stolon and more than two gastrozooids, (Fig.1D) and (4) dead polyps with detached or empty stolon.

### Testing survival of polyp colonies at low temperature

To test the ability of colonies to survive at low temperatures without feeding, we put Z4B colony in a 50 ml plastic ‘Falcon’ tube containing clean MFSW and kept at 4, 10 or 18°C for 4 weeks without feeding. The seawater was changed once after two weeks. Viability of the colonies was scored by the presence of polyps with tentacles, and further confirmed by recovery of the polyp colony after 15 days at 18°C with regular feeding.

### Testing of treatments to efficiently “sterilize” *Clytia* polyps and jellyfish

To avoid environmental release of genetically modified *Clytia* strains, we examined effective ‘sterilization’ by osmotic shock. Polyps, medusae and planula larvae were incubated in 1/10 diluted artificial seawater by adding 10 volumes of tap water to the culture container and incubated for different times (1 to 5 min for medusae and planula; 30 to 120 min for polyps). Complete loss of living tissue was evaluated under the microscope immediately and 24 hours after the treatment.

## Results

### Growth of *Clytia* jellyfish and polyp colonies

We first characterized *Clytia* Z4B female medusa growth using the standard culture set up (see supplementary protocol). *Artemia salina* n nauplii were used to feed twice a day (Fig. 4A). Medusae were grown in nursery tank until the average bell diameter reached 2.5 mm (dotted line 1). Then medusae were transferred to a Kreisel tank. Density was reduced to 50 medusae/tank (roughly 10 medusae/1liter) once they grew to 5.0 mm bell diameter (dotted line 2). The newly released jellyfish took 6 days to grow to 2.5 mm bell diameter. Once transferred to a Kreisel tank, the growth of the medusae accelerated, exhibiting a dual-phase growth curve (Fig.4A). The medusae reached adult size (10∼12 mm) 14 days after release, when the ovulation was also observed.

**Figure 4.**
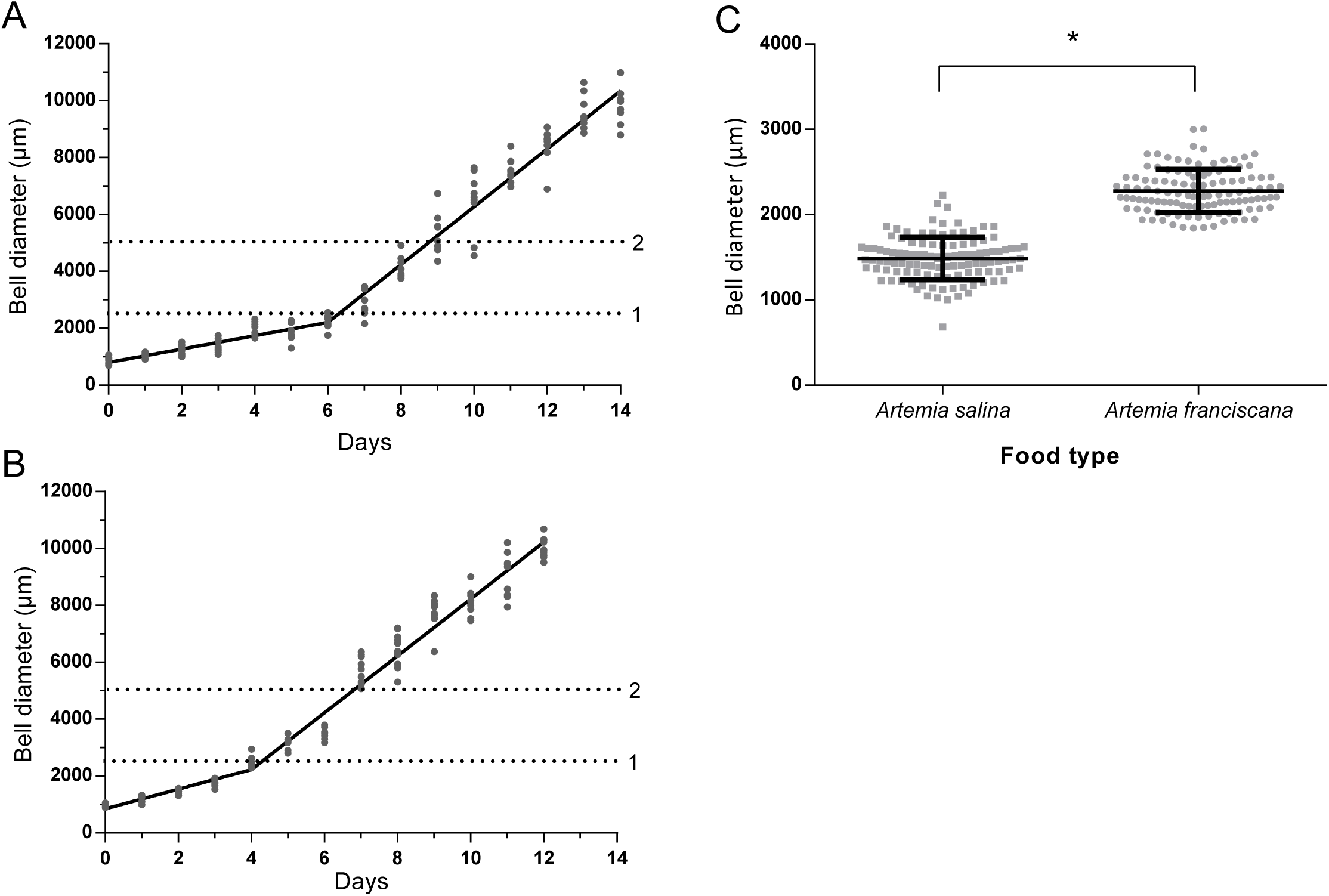
Growth of jellyfish. (A) Growth of jellyfish fed with *Artemia salina* (2-segment linear model, R^2^= 0.98) and (B) with *Artemia franciscana* (R^2^= 0.98). Dotted lines 1 and 2 represent 2.5 mm and 5 mm bell diameters, thresholds to transfer medusae from nursery tank to Kreisel tank and to decrease the density to 50 respectively. The segment boundary for 2-phase growth was defined by date to transfer medusae. (C) Size comparison of n=115 6 days-old jellyfish fed with *Artemia salina* and *Artemia franciscana* (t-test * p<0.05).

As the relatively slower growth observed over the first days spent in the nursery tank may be due to the lower capacity for food capture in the juvenile stage, we assessed different food types. We found that smaller *Artemia franciscana* nauplii larvae, hatched from cysts produced in Vietnam, achieved the best growth of juvenile stage medusae (Fig.4B). These A. *franciscana* nauplii are significantly smaller (body length 601 ± 107 µm at instar 3 larva stage) than standard *A. salina* nauplii (783 ± 97 µm, Fig.S2). By feeding with *A. franciscana* nauplii, young jellyfish reached 2.5 mm diameter in 4 days instead of 6 days for *A. salina* (Fig. 4B). We directly compared the early phase of medusae growth between groups fed with these two types of Artemia (Fig. 4C). We cultured 115 baby medusae/ tank and added equivalent volumes of *A. salina* and *A. franciscana* nauplii. The bell diameter of young medusa was significantly larger with *A. salina* 6 days later (Fig 4C). These results suggest that feeding efficiency is critical for juvenile medusae growth. On the other hand, for in later stage medusa, the growth speed was similar between two food types.

Unlike medusa stages, the polyp colony grows asexually (vegetatively) and can be considered practically immortal, providing ideal material for conserving genetic strains. We measured the growth speed of the polyp colony under daily feeding by counting the number of gastrozooids (feeding polyps) and gonozooids (medusa budding polyps) (Fig. 5). We made transplant colonies of identical Z4B strains (see methods). Starting from colonies with 5 gastrozooids, the colony size roughly doubled to more than 10 gastrozooids in 7 days and reached 30∼40 gastrozooids in 13 days (Fig. 5A). The colonies formed a first gonozooid within 4 days and acquired more than 10 gonozooids in two weeks (Fig. 5B). No clear difference in colony growth was observed between feeding once and twice per day. Feeding once per day is thus sufficient for the maintenance of wild-type colonies.

**Figure 5.**
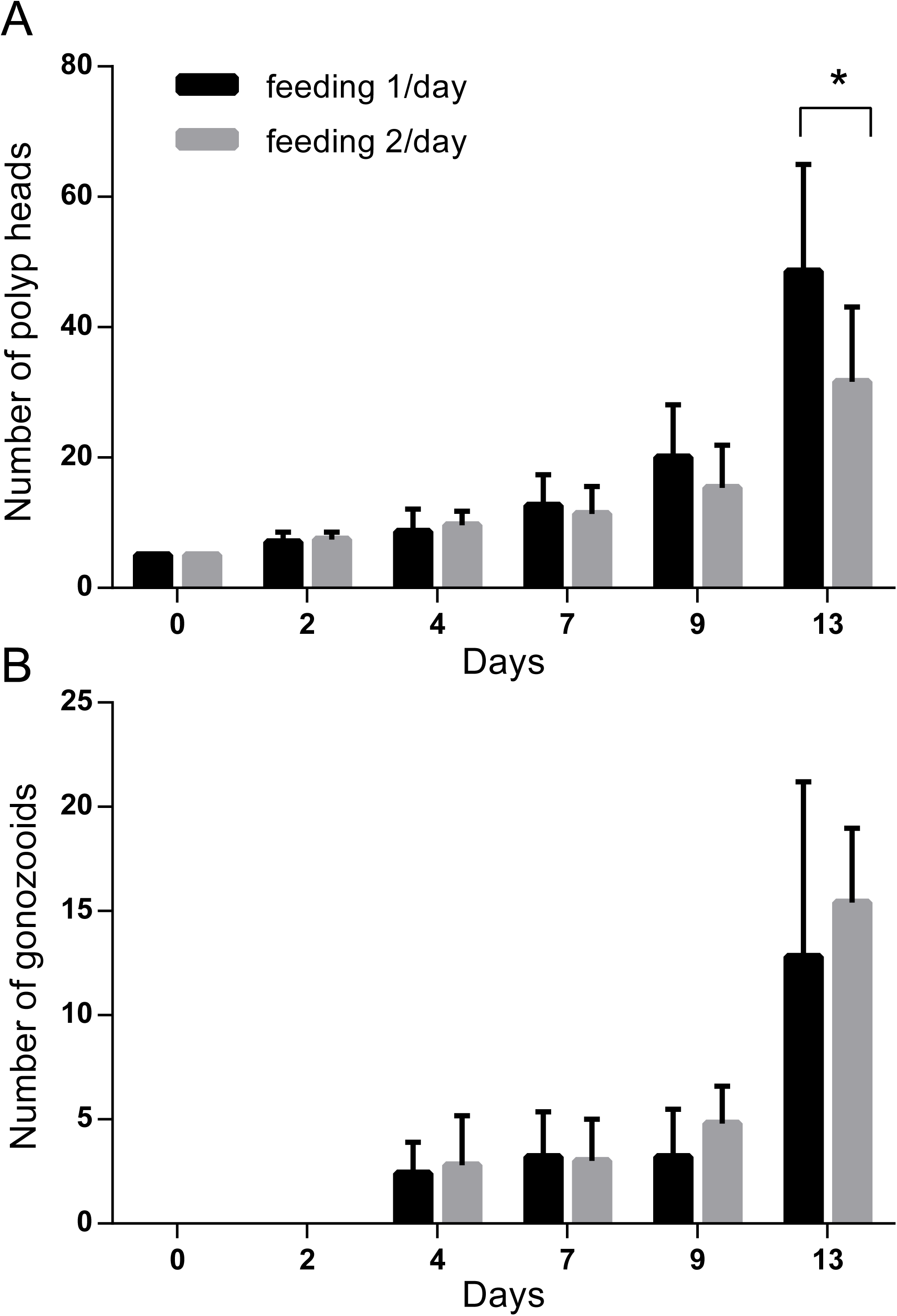
Growth of polyp colonies. (A) Polyp colony growth measured by number of polyps (gastrozooids) and (B) formation of gonozooid polyps with one or two feedings/day (two-way ANOVA * p<0.05).

### Metamorphosis of planula larvae

Once they have reached adult size with mature gonads, *Clytia* medusae can release eggs and sperm daily for several weeks following a dark-to-light cue, providing convenient material for studies of embryogenesis and larva formation, as well as for the creation of gene-edited *Clytia* lines. A critical step in generating mutant strains is to induce and successfully complete metamorphosis of planula larvae into polyp colonies. In natural conditions, metamorphosis is triggered by unknown cues from bacterial biofilms, and involves downstream cellular responses mediated by GLWamide family neuropeptides (Takahashi and Takeda, 2015). In the laboratory, a synthetic GLW-amide neuropeptide (GLWamide2, GNPPGLW-NH2) identified from the *Clytia* transcriptome sequences was used to trigger metamorphosis (Momose et al., 2018; Quiroga Artigas et al., 2018). The GLWamide2 sequence is similar to known hydrozoan metamorphosis-inducing peptides like metamorphosin A (EQPGLW-NH2) for *Hydractinia echinata* (Leitz and Lay, 1995) and KPPGLW-NH2 for *Phialidium (Clytia) gregarium* (Freeman, 2005). We found that metamorphosis efficiency of 3-day-old planulae generated from Z strain crosses on glass slides (Fig. 6) could reach only 40∼60%. This is considerably lower than the settlement efficiency reported for *Clytia gregarium* (Freeman, 2005). Neither the density of the planulae nor the use of plastic rather than glass substrates affected the efficiency, so we sought to identify more efficient GLWamides. Published *Clytia* neuropeptide precursor sequences (Takeda et al., 2018) include two pro-neuropeptide genes (*Che-pp2* and *Che-pp11*) that each encode several potential GLWamide precursors. We tested 15 chemically synthesized neuropeptides (Table 1) derived from these sequences to see whether any were more active that GLWamide2. While most of the variants were able to induce metamorphosis at 5 µM, none were efficient at 1 µM or less, except for GLWamide-6 (pyroGlu-QQAPKGLW-NH3). GLWamide-6 was active at as low as 0.3 µM, inducing virtually all planula to settle on the glass slide (Fig. 7). GLWamide-6 is thus so far, the most effective inducer of *Clytia* larval settlement and metamorphosis in vivo.

**Figure 6.**
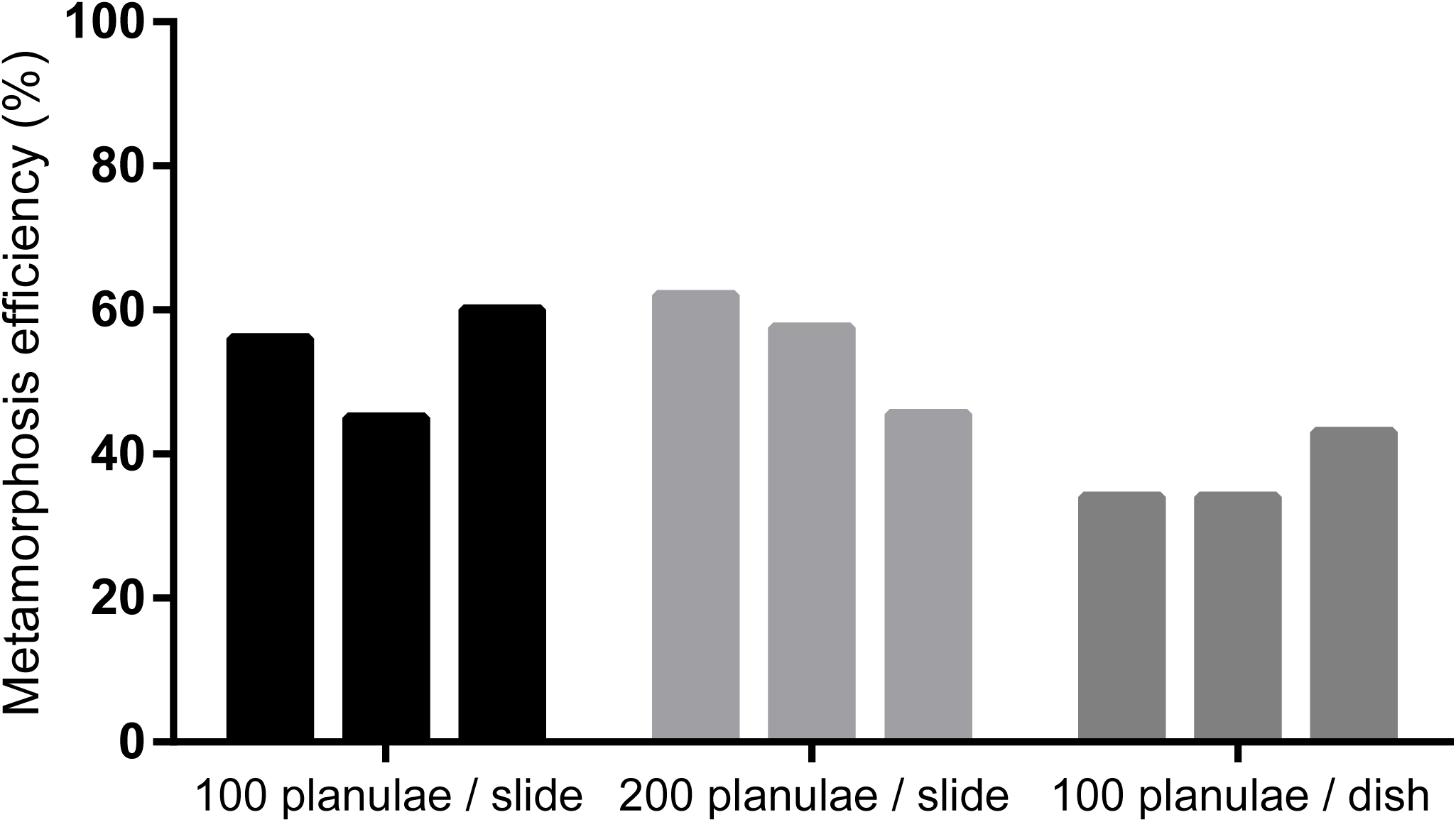
GLW-amide induced metamorphosis efficiency is not affected by substrate material or planula density. Metamorphosis of planulae induced by GLW-amide2. Efficiency was measured by settlement at different density or on different materials. 100 or 200 planulae were induced metamorphosis on 100 mm plastic petri dish and 75 × 50 mm glass slides, repeated three times for each experiment.

**Figure 7.**
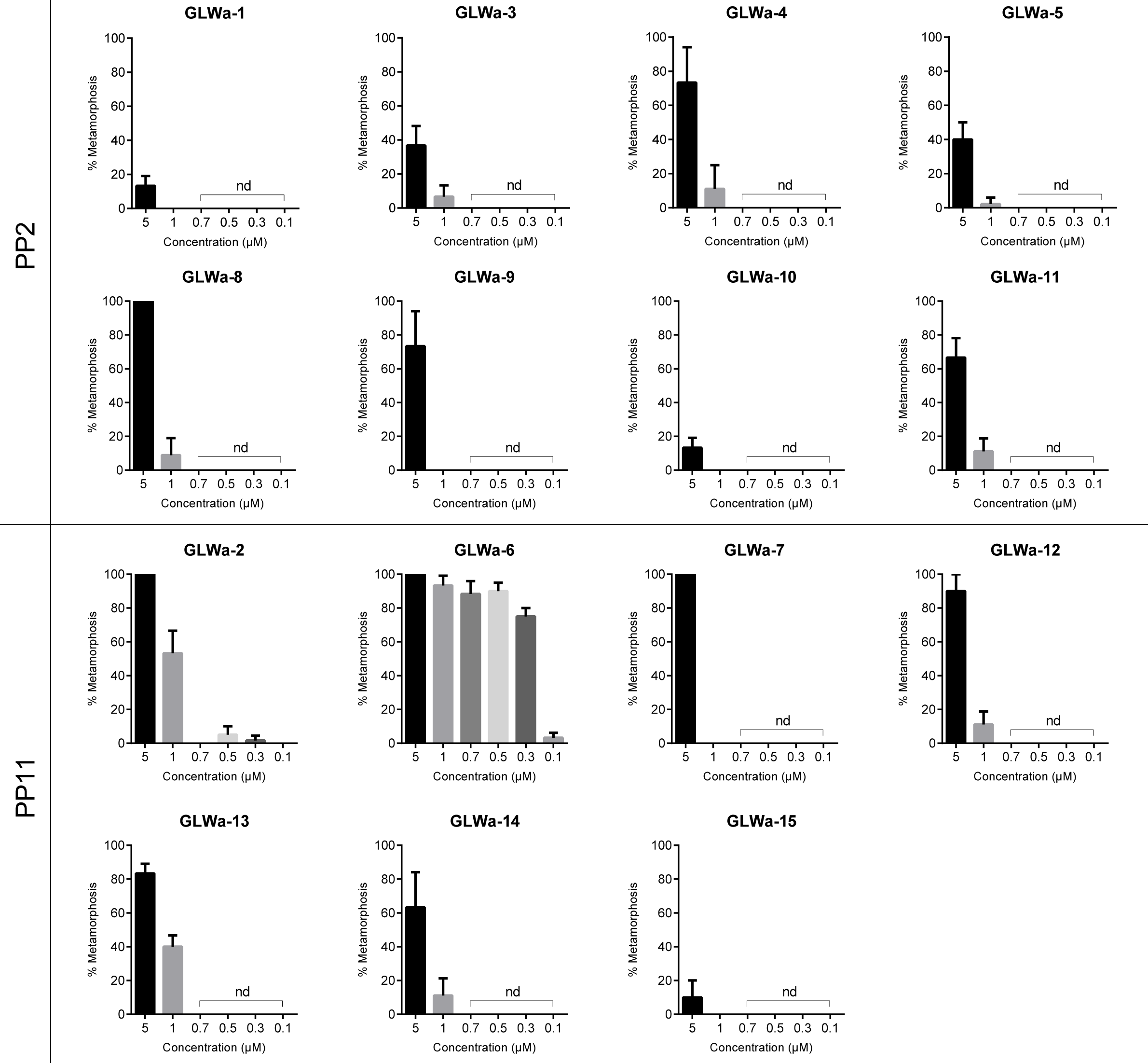
Metamorphosis efficiency with different synthetic GLW-neuropeptides. Synthetic GLWamide neuropeptides predicted from the ChePP1 and ChePP2 precursors were synthesized and their metamorphosis inducing efficiency was tested. The list of peptide sequence is in Table 1. Experiment was repeated for 5 times for each condition with 20 planulae.

**Table 1.** List of tested synthetic neuropeptides

| Name | Peptide | Precursor (Pro-neuropeptide) |
| --- | --- | --- |
| GLWa-1 | GAVQGLW-NH<sub>2</sub> | PP2 |
| GLWa-3 | pGlu-NSPGALGLW-NH<sub>2</sub> | PP2 |
| GLWa-4 | PGAVQGLW-NH<sub>2</sub> | PP2 |
| GLWa-5 | pGlu-GLW-NH<sub>2</sub> | PP2 |
| GLWa-8 | PGALGLW-NH<sub>2</sub> | PP2 |
| GLWa-9 | LGLW-NH<sub>2</sub> | PP2 |
| GLWa-10 | KPGAVQGLW-NH<sub>2</sub> | PP2 |
| GLWa-11 | SPGALGLW-NH<sub>2</sub> | PP2 |
| GLWa-2 | GNPPGLW-NH<sub>2</sub> | PP11 |
| GLWa-6 | pGlu-QQAPKGLW-NH<sub>2</sub> | PP11 |
| GLWa-7 | pGlu-PGNPPGLW-NH<sub>2</sub> | PP11 |
| GLWa-12 | pGlu-QPGNPPGLW-NH<sub>2</sub> | PP11 |
| GLWa-13 | NPPGLW-NH2 | PP11 |
| GLWa-14 | PPGLW-NH2 | PP11 |
| GLWa-15 | GPIPGMW-NH2 | PP11 |

**Table 2.**
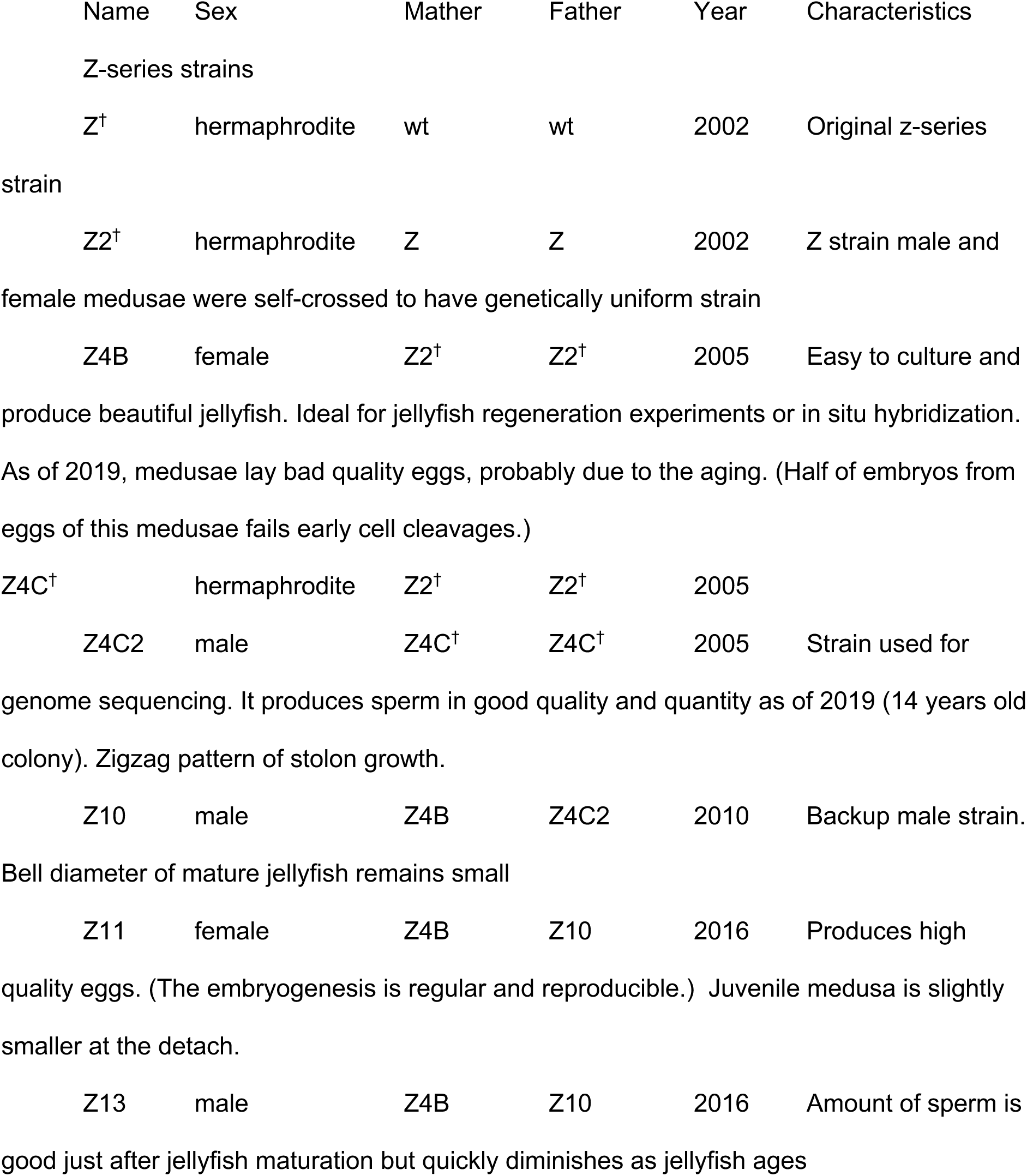
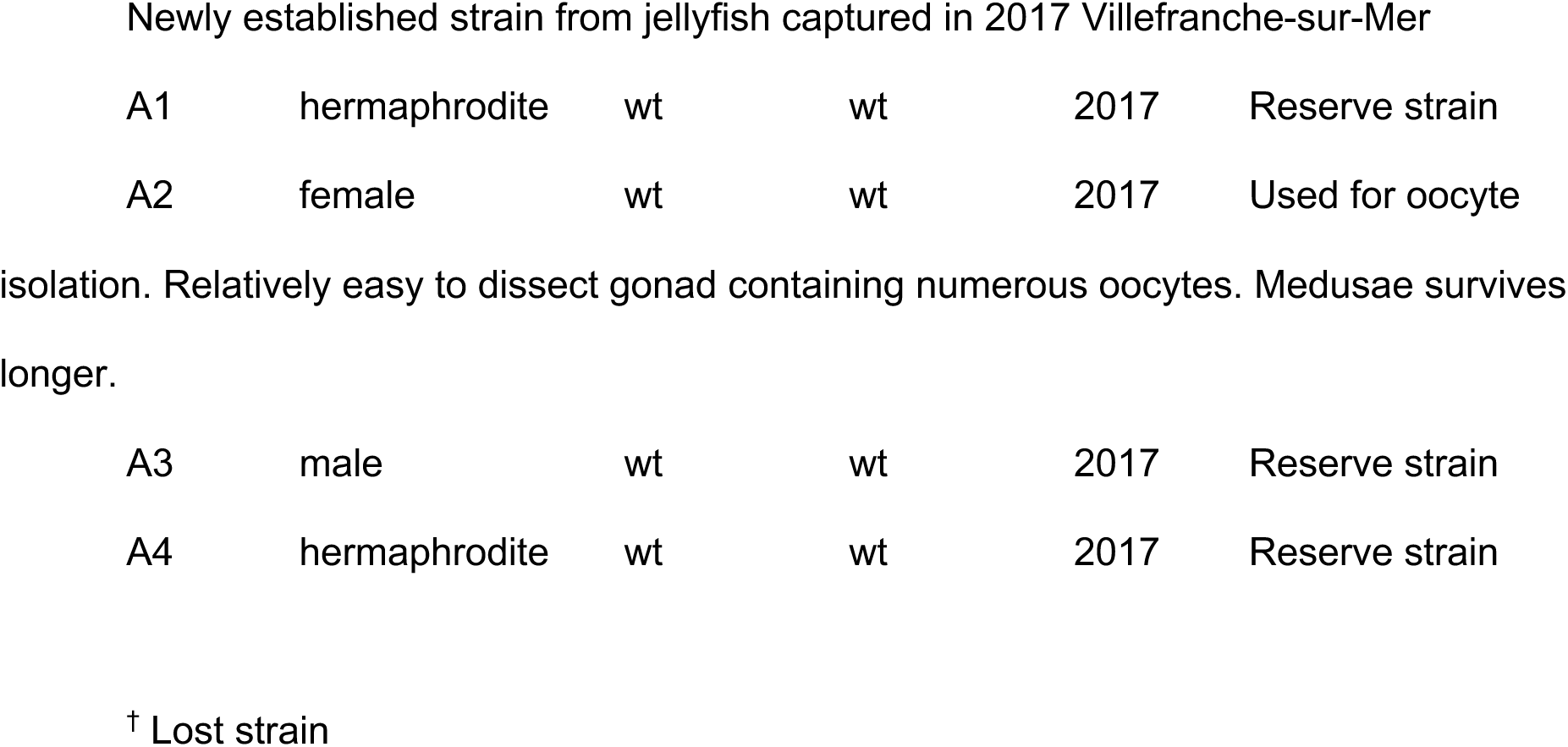
List of *Clytia* wildtype strains.

Survival and successful growth of the primarily polyp into a multi-polyp colony is another critical life cycle step. Only about 10% of successfully settled primary polyps formed a polyp colony in a few weeks (Fig. 8). This represents a significant bottleneck for mutant colony production of transgenic animals and mutants created for instance by CRISPR/Cas9 technology. As the size of the primary polyp is usually smaller than that of gastrozooids within a colony (Fig.1), we hypothesized that food capture may be again one of the bottlenecks. We thus compared primary colonies fed either with intact *Artemia salina* larvae or with smashed ones using a 25-gauge syringe needle just before feeding. The smashed food improved the colony formation rate at least temporary (Fig. 8A). Feeding with smashed *Artemia franciscana* or smashed *Artemia salina* gave similar results (Fig. 8B). Smashing artemia nauplii thus improves young polyp colony survival even further durable feeding to the early stage polyp colonies will be needed.

**Figure 8.**
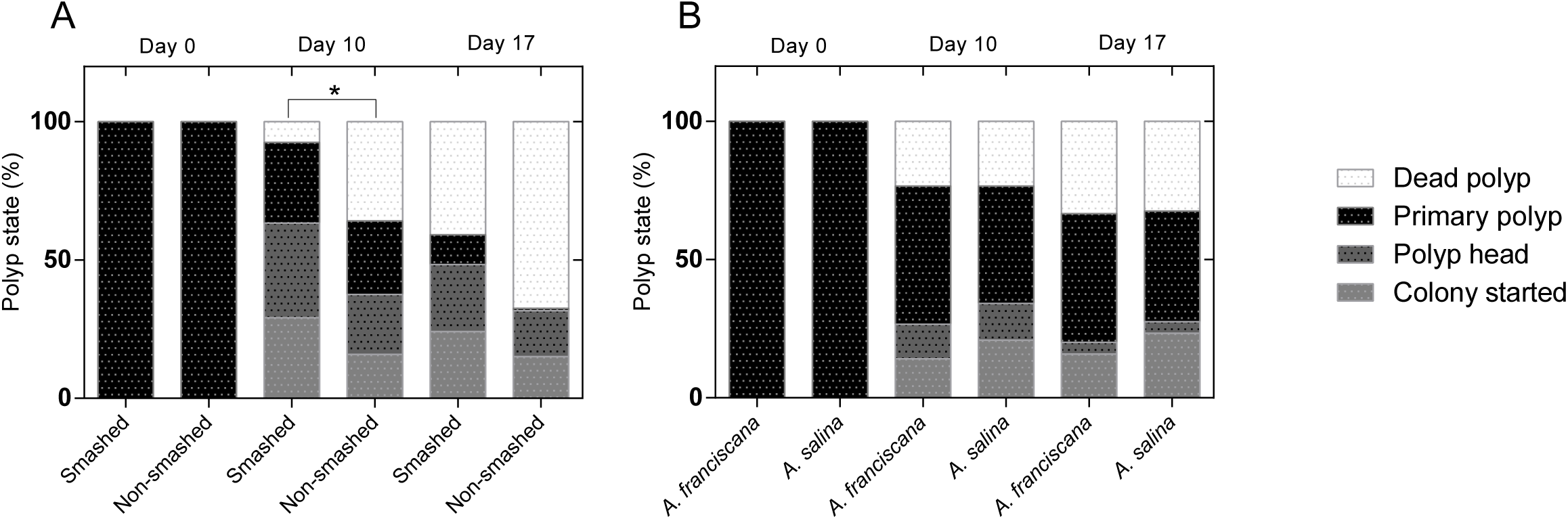
Primary polyp survival and colony formation efficiency. Survival, primary polyp formation (polyp head) and successful colony formation under different feeding regimes were measured for 40 successfully settled planula larvae (N=5 for each condition). (A) comparison of feeding with smashed and non-smashed artemia nauplii, showing improved colony formation using smashed artemia (two-way ANOVA, * p<0.05). (B) comparison of smashed Artemia franciscana and smashed *Artemia salina* showing no clear difference.

### Storage of polyp colonies

Vegetatively growing, regularly fed, *Clytia* polyp colonies can be maintained for a long period of time without a genetic cross. Some hydrozoan species, such as *Cladonema pacificum*, can be maintained at 4°C for several months to years in a dormancy state (Ryusaku Deguchi personal communication). Developing such colony dormancy methods in *Clytia* would allow maintaining genetic strains for long periods with minimal care. We found that after one month of storage at 4°C, *Clytia* colonies lose polyps and do not recover when cultured at 18°C. In contrast, functional gastrozooids survived in all colonies stored at 10°C and were able to eat and grow immediately after transferring to 18°C, indicating that *Clytia* colonies are resistant against starvation when maintained at low temperatures (Fig.S2).

### Sterilization

To avoid any risk of contamination of laboratory strains into the natural environment, it is important to eliminate any living material before discarding. We found that an effective method to kill *Clytia* planulae, polyps and medusae is exposure to low salinity water. Medusae and larvae disintegrated completely within 5 minutes when soaked in low osmolarity water (3.7‰; x1/10 salinity, Fig. S3A). The presence of the protective theca renders the polyp colony stage more resistant but 1 hour of incubation was sufficient to fully dissolve polyp colonies (Fig. S3B).

## Discussion

We have developed a culture system for *Clytia hemisphaerica*, one of the hydrozoan model species suitable for developmental biology. The presence of a medusa stage in the *Clytia* life cycle opens up many research possibilities. Its transparent body is suitable for cellular studies, including fluorescent microscopy. Medusae mature rapidly and spawn gametes once a day, reliably induced by a simple dark-light cue (Quiroga Artigas et al., 2018). In this work we described an optimized *Clytia* whole life cycle culture system developed so that biologists can reliably maintain and produce animal resources in the laboratory. The shortest full sexual generation cycle time for genetics studies in this system is about 2 months. A genetic strain can be maintained for many years as a polyp colony and be shared by making colony duplicates.

### Medusa growth: the importance of early stage feeding

In our experiences, a colony established on a large (75 × 50 mm) slide maintained in good nutritional conditions will typically produces several dozen of baby medusae each day. The measurements reported here revealed that the medusae grows to adult size and sexual maturity (about 10 mm in diameter) in less than two weeks – a time course suitable for genetic studies. The first few days, i.e.until the juvenile medusae reaches 2.5 mm bell diameter, were found to be critical for successful jellyfish growth for two reasons. Firstly, these small medusae cannot be accommodated in the Kreisel tanks as they pass through the nylon retaining filters. Closed tanks such as bubble-circulating nursery tanks or crystallizing dishes can be used for this stage, though require additional attention to the water quality with overfeeding being particularly harmful. Alternative systems for the early step of medusa culture could be developed in the future, for instance to avoid the harmful effect of air bubbles that can damage medusae as they reach larger sizes. Secondly, juveniles cannot efficiently catch live *Artemia salina* nauplii. This issue was mitigated by using the smaller nauplii of *A. franciscana*, which significantly improved the growth of baby medusae and shorted the time to adulthood. Inefficient feeding may potentially be problematic for some mutant strains, for instance with developmental or neuronal phenotypes. Direct delivery of food to medusa mouth (“hand feeding”) may be required in this case.

### Maintaining the polyp colony

Another advantage of *Clytia* as a genetic model animal is the robustness of the polyp colony stage. A colony can rapidly extend stolons and generate new polyps in a vegetative manner. Commonly available *Artemia* nauplii are convenient and sufficient as food. Our tests showed that the colony size can double in less than one week, and that feeding once a day is sufficient at least for wild type strains. Colony growth does not seem to be limited by age or number of polyps. For example, the Z4B strain used in this work was established in 2005 and still propagates rapidly (Leclère et al., 2019). The polyp colony stage is thus useful for long-term maintenance of genetically modified strains created by CRISPR/Cas9 technology.

The standard aquaculture system described here use artificial sea water (37‰) adjusted to the salinity of the Mediterranean water of the bay of Villefranche-sur-Mer where our *Clytia* founder animals were collected (http://www.obs-vlfr.fr/data/view/radehydro/std/; Leclère et al 2019). The species *Clytia hemisphaerica* is present worldwide and some local populations can certainly tolerate lower salinity, so colonies from animals collected from other localities might prefer other salinities. Medusae seems to be affected when the salinity reaches to 39‰ by evaporation. On the other hand, polyps from our laboratory *Clytia* strains was not largely affected when by accident the salinity increased up to 42‰ for a few days in our laboratory. Though it is highly recommended to maintain the salinity within from 37‰ to 38‰. Long-term physiological effect of the extreme salinity is to be examined. We maintain most of the colonies and jellyfish at about 18∼20°C. Using systems maintained at 24°C during the early stages of polyp colony growth seems to favor irreversible determination to female. Medusa sex is partly determined by colony temperature according to the studies using hermaphroditic strains. (Carré and Carré, 2000). On Temperatures higher than 24°C seem harmful for polyp colony survival (unpublished observations). At lower temperatures, some hydrozoan species undergo dormancy making dormant structures such as podocysts or dormant coenosarc (stolon) (Calder, 1990; Thein et al., 2012). *Clytia* colonies could survive for one month of starvation when they were kept at 10°C retaining smaller but functional polyps, while feeding was necessary to maintain polyps at 18°C. Considering the lowest winter sea water temperature (12°C at 1 m and 50 m depth) where the original *Clytia* jellyfish are collected (http://www.obs-vlfr.fr/data/view/radehydro/std/), it is likely to be a hibernation state. Further tests will be necessary to develop reliable long-term methods for strain maintenance including cryoconservation.

The *Clytia* polyp colony can grow by constantly extending stolons and increasing the number of polyps with an appropriate food supply. Constant growth by new stolon extension is critical to maintain the colonies because individual gastrozooids are not immortal, unlike a whole colony. Colony growth is sensitive to various factors. For instance, stolon extension can be blocked by algae covering the glass surface. It is thus necessary to remove the old part of the colony and clean glass surface covered by algae by wood toothpick or plastic scraper every once in a few months. Reducing the luminosity or temperature of the culture environment prevents the growth of red algae. Further, occasional flushing of culture plates by pipetting will also help to reduce accumulation of the organic particles such as food leftovers. Making duplicates of genetic strains by polyp transplantation (see materials and methods) provides a convenient way to share genetic strains in the community. Finally, *Clytia* polyps, which are potentially invasive given their extremely high capacity for vegetative growth, can be easily eliminated within one hour of adding ten volumes of tap water, when a part of the colony is detached from the culture plates. Other physical or chemical confinement methods such as UV-C irradiation or sodium hypochlorite treatment could also employed in transgenic *Clytia* facilities.

### From embryos to polyp colonies

Once they become sexually mature (2-3 weeks after budding from the gonozooid depending on feeding), adult medusae daily spawn eggs or sperm, depending on the sex, for several weeks. After fertilization, embryos develop to form simple ciliated planula larvae with an elongated “oral-aboral” body axis. By using the newly designed synthetic neuropeptide GLWamide6 we were able to increase the planula settlement efficiency to nearly 100%. There is still room for improvement concerning metamorphosis efficiency: with our current protocols only 25% of settled primary polyps survive and make colonies with multiple polyps (Fig.8). In most cases, settled planula fail to form functional gastrozooids. In other cases, a primary polyp forms but fails to extend a stolon or make additional polyps. This latter problem can be partly remedied by providing smashed and immobilized *Artemia* nauplii, which are easier for the primary polyps to trap and ingest. The effect of smashed *Artemia* is not significant enough in long term (17 days), suggesting that feeding to primary polyps needs further improvement.

## Supporting information

Supplemental Protocol

## Acknowledgments

This work benefited from the support of the project i-MMEJ (ANR-17-CE13-0016) of the French National Research Agency (ANR) and ASSEMBLE-Plus Joint Research Activity (JRA3). BW is a Howards Hughes Medical Institute Fellow of the Life Sciences Research Foundation. The CRBM is supported by EMBRC-France, whose French state funds are managed by the ANR within the Investments of the Future program under reference ANR-10-INBS-02. We thank Maciej Mańko for making new A-series wildtype strains during his stay in Villefranche-sur-Mer. *Clytia hemisphaerica* wildtype strains used in this work are available from CRBM (https://www.embrc-france.fr/fr)

## References

Carré, D., and Carré, C. (2000). Origin of germ cells, sex determination, and sex inversion in medusae of the genus Clytia (Hydrozoa, leptomedusae): the influence of temperature. J. Exp. Zool. 287, 233–242.

Cook, C.E., Chenevert, J., Larsson, T.A., Arendt, D., Houliston, E., and Lénárt, P. (2016). Old knowledge and new technologies allow rapid development of model organisms. Mol. Biol. Cell 27, 882–887.

Freeman, G. (2005). The effect of larval age on developmental changes in the polyp prepattern of a hydrozoan planula. Zoology (Jena) 108, 55–73.

Calder, D.R. (1990). Seasonal cycles of activity and inactivity in some hydroids from Virginia and South Carolina, U.S.A. Can. J. Zool. 68, 442–450.

Galliot, B. (2012). Hydra, a fruitful model system for 270 years. Int J Dev Biol 56, 411–423.

Gold, D.A., Katsuki, T., Li, Y., Yan, X., Regulski, M., Ibberson, D., Holstein, T., Steele, R.E., Jacobs, D.K., and Greenspan, R.J. (2019). The genome of the jellyfish Aurelia and the evolution of animal complexity. Nat Ecol Evol 3, 96–104.

Goldstein, B., and King, N. (2016). The Future of Cell Biology: Emerging Model Organisms. Trends Cell Biol 26, 818–824.

Greve, W. (1968). The “Plankton Kreisel” a new device for culturing zooplankton. Marine Biology 1, 201–203.

Houliston, E., Momose, T., and Manuel, M. (2010). Clytia hemisphaerica: a jellyfish cousin joins the laboratory. Trends Genet 26, 159–167.

Leclère, L., Copley, R.R., Momose, T., and Houliston, E. (2016). Hydrozoan insights in animal development and evolution. 39, 157–167.

Leclère, L., Horin, C., Chevalier, S., Lapébie, P., Dru, P., Peron, S., Jager, M., Condamine, T., Pottin, K., Romano, S., et al. (2019). The genome of the jellyfish Clytia hemisphaerica and the evolution of the cnidarian life-cycle. Nat Ecol Evol 3, 801–810.

Leitz, T., and Lay, M. (1995). Metamorphosin A is a neuropeptide. Rouxs Arch. Dev. Biol. 204, 276–279.

Lilley, M.K.S., Ferraris, M., Elineau, A., Berline, L., Cuvilliers, P., Gilletta, L., Thiery, A., Gorsky, G., and Lombard, F. (2014). Culture and growth of the jellyfish Pelagia noctiluca in the laboratory. Marine Ecology Progress Series 510, 265–273.

Momose, T., and Concordet, J.-P. (2016). Diving into marine genomics with CRISPR/Cas9 systems. Mar Genomics 30, 55–65.

Momose, T., and Houliston, E. (2007). Two oppositely localised frizzled RNAs as axis determinants in a cnidarian embryo. PLoS Biol 5, e70.

Momose, T., De Cian, A., Shiba, K., Inaba, K., Giovannangeli, C., and Concordet, J.-P. (2018). High doses of CRISPR/Cas9 ribonucleoprotein efficiently induce gene knockout with low mosaicism in the hydrozoan Clytia hemisphaerica through microhomology-mediated deletion. Sci Rep 8, 11734.

Plickert, G., Frank, U., and Müller, W.A. (2012). Hydractinia, a pioneering model for stem cell biology and reprogramming somatic cells to pluripotency. Int J Dev Biol 56, 519–534.

Quiroga Artigas, G., Lapébie, P., Leclère, L., Takeda, N., Deguchi, R., Jékely, G., Momose, T., and Houliston, E. (2018). A gonad-expressed opsin mediates light-induced spawning in the jellyfish Clytia. Elife 7, 81.

Ramondenc, S., Ferrieux, M., Collet, S., Benedetti, F., Guidi, L., and Lombard, F. (2019). From egg to maturity: a closed system for complete life cycle studies of the holopelagic jellyfish Pelagia noctiluca. J. Plankton Res. 41, 207–217.

Raskoff, K.A., Sommer, F.A., Hamner, W.M., and Cross, K.M. (2016). Collection and Culture Techniques for Gelatinous Zooplankton. The Biological Bulletin 204, 68–80.

Spangenberg, D.B. (1965). Cultivation of the life stages of Aurelia aurita under controlled conditions. J. Exp. Zool 159, 303–318.

Takahashi, T., and Takeda, N. (2015). Insight into the molecular and functional diversity of cnidarian neuropeptides. Int J Mol Sci 16, 2610–2625.

Takeda, N., Kon, Y., Quiroga Artigas, G., Lapébie, P., Barreau, C., Koizumi, O., Kishimoto, T., Tachibana, K., Houliston, E., and Deguchi, R. (2018). Identification of jellyfish neuropeptides that act directly as oocyte maturation-inducing hormones. Development 145, dev156786.

Thein, H., Ikeda, H., and Uye, S.-I. (2012). The potential role of podocysts in perpetuation of the common jellyfish Aurelia aurita s.l. (Cnidaria: Scyphozoa) in anthropogenically perturbed coastal waters. In Jellyfish Blooms IV, (Dordrecht: Springer, Dordrecht), pp. 157–167.

