## Supplemental Protocol for "An improved whole life cycle culture protocol for the hydrozoan genetic model *Clytia hemisphaerica*"

### *Clytia hemisphaerica* culture protocol

#### Tools and glass/plasticware.

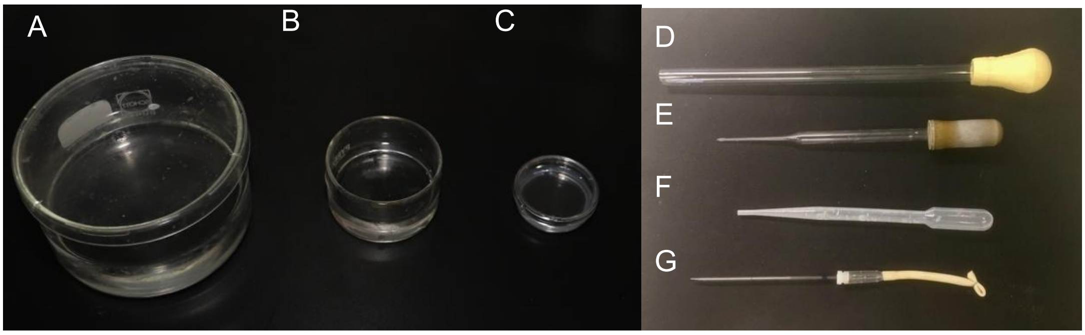


Figure 1. glass/plastic wares. See the main text for the detail.

- Crystallizing dish
  - Large: φ 95 mm, height 5~6 cm (Fig.1A)
  - Small: φ 50 mm, height 3 cm (Fig.1B)
- Petri dish: φ 35 or 50 mm (Fig.1C)
- Jellyfish transfer pipette: φ(ext) 10 mm glass or plastic pipes with pipette bulb for transferring medusae between tanks/dishes (Fig.1D)
- PASTEUR pipette: for feeding (Fig.1E)
- Transfer pipette (3~5 ml size for cell culture, non-sterile, ex. Samco Scientific 3 ml “225”: for feeding, fertilization (sperm transfer) or small medusae less than 5 mm Fig.1F)
- Embryo/egg transfer pipette; made by pulling glass micropipette (ex Hirschmann ringcaps 100 µl, cat no. 9600199) under alcohol lamp flame. The tip is cut with a diamond pen and flame rounded. Used with a pipette bulb (ex. Hirschmann cat no 9650101) Fig.1G
- Large size glass slides (75 mm x 50 mm)
- Slide staining jar with lid (Kartell 235305) and basket (Kartell 235405): for holding and transferring slides.
- Diamond pen (to label slides)
- Petri dish (φ 100 mm, for the rid of crystalizing dish and keeping glass slides during metamorphosis)
- Glass plates (160 mm x 140 mm) for large scale production of medusae
- microscissors (Corneal scissor, ex. Moria surgical, 8100)

##### 3D printed materials

- Single slide holder (https://pinshape.com/items/35213-3d-printed-glass-slide-holder-for-aquarium )
- Glass plate holder (https://pinshape.com/items/35215-3d-printed-glass-plate-holder-for-hydrozoan-polyp-culture )
- Kreisel tank filter kit (v.3) (https://pinshape.com/items/52438-3d-printed-kreisel-tank-filter-kit-ver-3 )

### Sea water preparation

- Reverse-osmosis (RO) water (eg produced using the General Electric, Merlin reverse osmosis system for drinking water)
- RedSea salt (https://www.redseafish.com/red-sea-salts/ )
- Plastic (polypropylene) tank
- Submersible water pump (ex. Eheim CompactOn 300)
- Digital refractometer (ex. Atago PR-100SA)

We use artificial seawater with salinity adjusted to 37‰ for all culture steps. Sea water with 37‰ salinity can be made by dissolving 40 grams of RedSea salt mixture is dissolved in 1 liter RO water.

##### Making 50 liters of artificial sea water

1. In the tank add 2.0 kg of RedSea Salt and to 48~ liter of RO water (use pre-labeled level mark).
2. Dissolve by mixing with a submersible water pump for several hours to one night.
3. Measure the salinity with digital refractometer and adjust it to 37‰ by adding RO water

##### Millipore filtered sea water (MFSW)

Filter the artificial sea water with 0.22 µm Millipore filter for embryo/planula culture and metamorphosis.

Notes:

We use RO water to remove chlorine in tap water. A restaurant grade RO water system is sufficient for this purpose. Alternatively use sodium thiosulfate to remove chlorine.

Of artificial sea water brands tested, RedSea salt (RedSea) and SeaLife (Marine Tech, Japan), both made at least partly from natural sea water, worked well. Artificial seawater reconstituted by purified salts (so called synthetic sea salt, for example Instant Ocean) affected spawning.

We have not systematically tested the effects salinity changes, but, our *Clytia* cultures are resistant to salinity changes within the range of 35‰ to 42‰ for at least several days. It is, however, recommended to regularity check and adjust the salinity to 37‰ by adding RO water in the reservoir, for the water-circulating system.

### Temperature control and sex determination.

Our standard culture temperature is 18 - 20°C. *Clytia* medusae grow well between 15°C and 21°C. At 25°C medusae grow poorly (Matsakis, 1993). Polyp colonies show a similar temperature preference. At 24°C, colony growth is significantly slower.

While the standard temperature is recommended for routine culture of polyp colonies, culture at higher (24°C) temperatures for several weeks or more from primary polyp stage may be used to favor the establishment of female strains,

Note: The sex determination seems affected by the temperature of polyp and medusa culture according to work using hermaphrodite *Clytia* colonies established from wild type medusae acquired in Villefranche-sur-Mer (Carré and Carré, 2000). A hermaphrodite colony produces both male and female medusae, and the population is affected by the temperature. In our recent experiences, however, hermaphrodite colonies are relatively minor, and most of the colonies produce exclusively male or female medusae. And the sex

The sex determination still seems epigenetically influenced by the temperature. The polyp colonies reared at 24°C tend to become female colonies and those grown at 18°C male.

The epigenetic sex determination seems to take place in early polyp colony stage, which is determinative and irreversible; only female medusae are produced from female colonies and male medusae from male colonies. The sex determination step, most likely during the initial polyp colony growth from primary polyp, still seems influenced by the temperature.

#### Configuration of seawater-circulating aquarium system

We designed culture tanks and the water circulation system to best suit the stage and size of jellyfish, and experimental needs (Fig.1 and Fig.2 in main text). Most of the jellyfish and polyp stages are maintained in a closed seawater recirculating aquaculture system. Each water-circulating aquarium unit was constructed on a scaffold of salt-resistant aluminum or plastic shelving equipped with a few dozen individual tanks on the upper shelves and a sea water reservoir (50~100 liter) on the bottom shelf.

##### Water circulation.

Sea water is delivered to each tank from the reservoir using a submersible pump (Eheim universal pump 3400). Overflow sea water is recovered by a drain pipeline and collected into a pre-chamber of the reservoir, where a nylon mesh filter (mesh opening size 200 µm) or a filter pad (Combo filter pad classic bonded & 50 micron fine water polishing filter, Aquatic Experts), and a protein skimmer (H&S aquaristik, A150-F2001) are installed to eliminate excess artemia, waste from *Clytia* as well as excess baby medusae released from polyp colonies. Treated seawater flows back into the main reservoir.

#### Thermostat and Lighting.

A water cooling/heating system (TECO TK-500 with submergible pump Eheim CompactOn 300) installed in the reservoir maintains sea water temperature with minimal temperature drift (TΔ=0.5°C). Walls and floors of the shelving were covered by black PVC plates (5 mm) and water-proof LED lighting was installed on the wall behind the tank to allow the transparent *Clytia* jellyfish to be visible by pseudo-dark-field illumination. In order to limit growth of red algae in the culture system, light exposure in the tanks was minimized, with LED lighting being wired to shut off automatically after use.

#### Salinity monitoring and adjustment

Salinity is measured by refractometer every few days and adjusted to 37‰ by adding RO water to the reservoir.

#### Tanks

Figure 2. Polyp tanks (left 3.5liter and 1.1liter) and Jellyfish "Kreisel" tank (right).

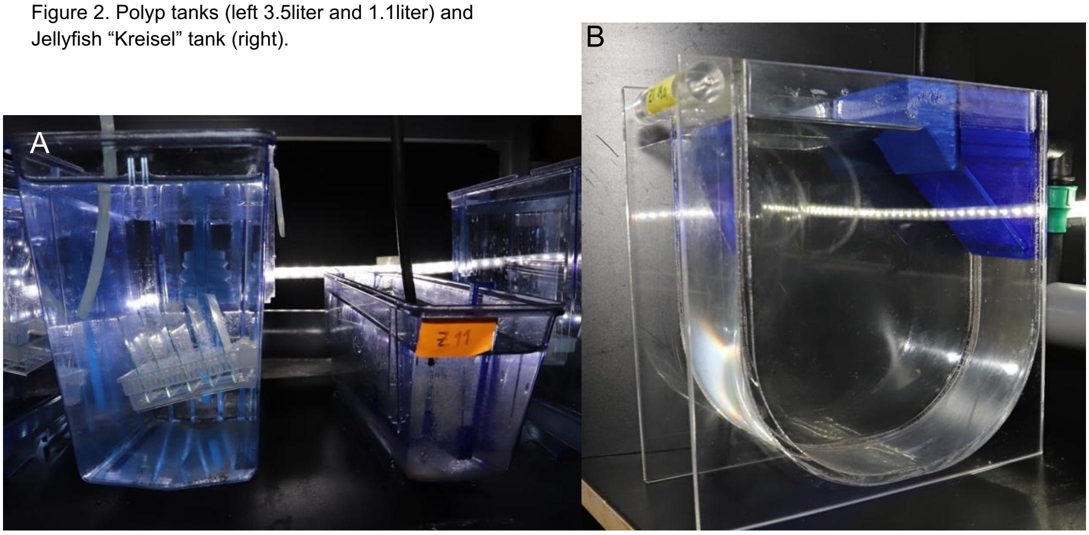


##### Polyp tanks

For polyp stages we use commercial zebrafish tanks (Fig. 1F, 1.1 liter model ZB11CP or 3.5 liter model ZB30CP, Tecniplast). Polyp colonies growing on glass slides (standard: 75 mm x 25 mm or wide: 75 mm x 50 mm) or larger glass plates ( 80 mm x 60 mm or 160 mm x 140 mm) are suspended on the side or the tank in histology slide baskets (Kartell, 20-Slide Microscope Slide Staining Dish, 235405) or 3D-printed plate holders (https://pinshape.com/items/35213-3d-printed-glass-slide-holder-for-aquarium).  Sea water is supplied continuously to each tank at a rate of 0.5~1.0 liter/min. Prior to the first use, polycarbonate plastic tanks need to be extensively cleaned and/or equilibrated by long soaking in sea water (up to 3 months) to remove residual chemical contamination.

##### Kreisel tanks

We culture adult medusae (over 2.5 mm size, typically from 6-days after release) in a modified version of the Kreisel tank (Fig.3) described by Greve (Greve, 1968; Purcell et al., 2013), simplified and optimized to maintain *Clytia* medusae healthily. The main tank was built from Poly(methyl methacrylate) (PMMA) plates (Fig.3A). Water nozzle and filter parts were made by 3D printing using polyethylene filament (Fig.3B, Volumic3D, PET-G Ultra filaments, 3D model available https://pinshape.com/items/52438-3d-printed-kreisel-tank-filter-kit-ver-3 ). These two parts are assembled using low-cost garden watering pipe connectors (LeroyMerlin, Aquaflow D1650 and D1005, see Fig.2B). It is critical to maintain a water speed of 10~20 mm/second near the tank periphery, obtained by adjusting the flow rate to 150~300 ml/min, depending on the opening diameter of the nozzle (1.5~ 2 mm x 4 nozzles). The key features of the Kreisel tank are; 1) to create circular water flow within the tank to keep medusae suspended and swimming, and 2) to filter particles such as dead Artemia or wastes by circulating seawater and filtering it.

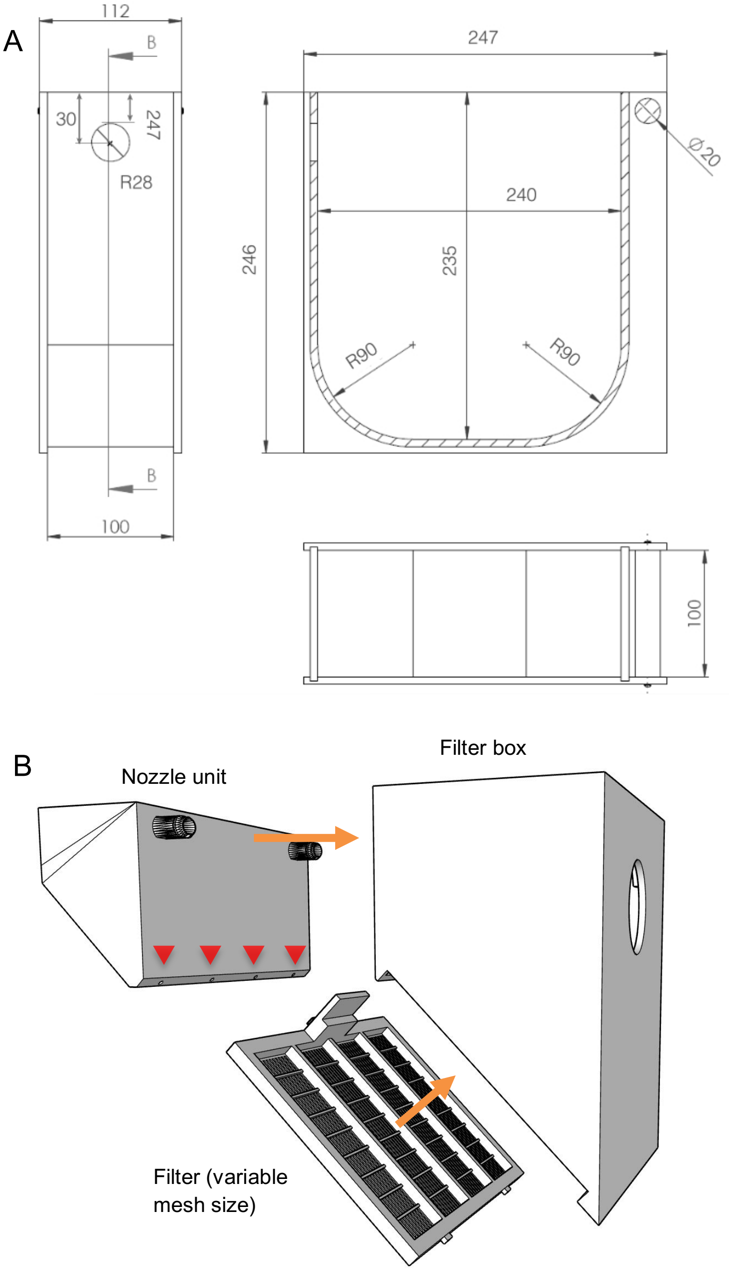


Figure 3. (A) An example of Kreisel tank for 50 adult medusae, to be combined with (B) 3D-printed filter/nozzle parts. These components are attached using connectors for garden watering pipes and an O-ring (Fig.2B). Water is supplied into the nozzle unit and comes out from nozzles (red arrowheads) located next to the filter unit. It makes rotary water flow and also prevent jellyfish form being trapped on the filter.

There are other types of Kreisel implementations (Purcell et al., 2013). For example, cylindrical walls installed in a commercial plastic tank may be used as a part of seawater-circulating system, where water flow is created by geared motor (see standalone beaker system described below) while slow water-circulation filters seawater.

Typically, *Clytia* medusae 2.5 mm diameter (about 5~7 days after budding) or larger can be hosted in our Kreisel tanks and can be maintained up to 5~6 weeks until their natural death. Up to 200 small (2.5~5 mm bell diameter) jellyfish may be maintained in one tank equipped with a fine mesh filter (200 μm mesh size). Once they have grown to 5 mm bell diameter or more, they are redistributed into multiple Kreisel tanks with coarse mesh filters (1 mm mesh pitch) at 40~50 jellyfish/tank.

#### Nursery tanks and crystallization dishes for juvenile medusa culture

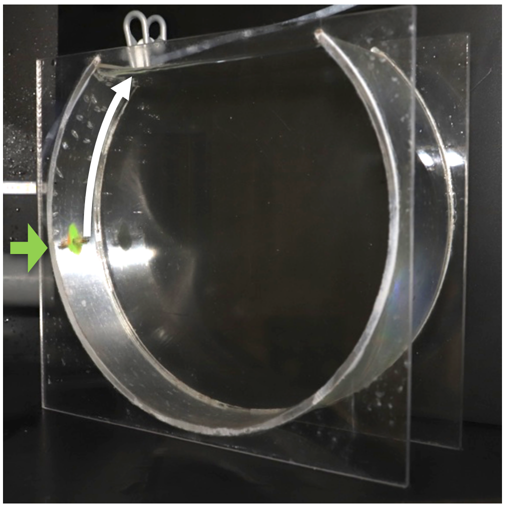

Figure 4. “Nursery” tank for collection, and hatching of young medusae less than 2.5 mm. The dram diameter: 270 mm (cutting 300 mm PMMA pipe longitudinally). Dram thickness: 80 mm. A φ5 mm (φint 3mm) PMMA air nozzle pipe is inserted on the side of the cylindrical wall (green arrow). The travel length of the bubble (white arrow), from the nozzle to water surface, needs to be long enough for the efficient rotation. The width of the opening is thus less than 170 mm. Similar tank is available from Schuran Kunststoffe (Breeding Air 300)

For some culture needs, notably for young medusae and embryos, we used individual tanks and dishes. A drum-shape "nursery" tank is used to collect and grow juvenile medusae up to 2 mm diameter (typically less than 6 days old after budding). Vertical water circulation is created by air bubbles (100~200 ml air/minutes) from 5 mm tubing inserted at one side of the shell, supplied by an air pump (ex. Eheim air pump 100). Larger medusae are more sensitive to turbulence created by bubbles and easily damaged in this system. Juvenile medusae need to be transferred to the Kreisel tank as soon as the average bell diameter reaches 2.5 mm in diameter.

Baby medusae can alternatively be collected and raised in 100 mm crystallization dishes, this method being preferable if the number or medusae is less than 100. These dishes are kept on a rotary shaker set at 50~70 rpm.  The sea water in the dish should be changed within a few hours after feeding. Equivalent dishes are also used to temporary concentrated adult jellyfish during gamete collection for short periods (typically less than 2 hours) during spawning. For longer culture no more than 5 or 6 adult medusae should be maintained in a dish.

#### Standalone beaker system (medusa and polyps)

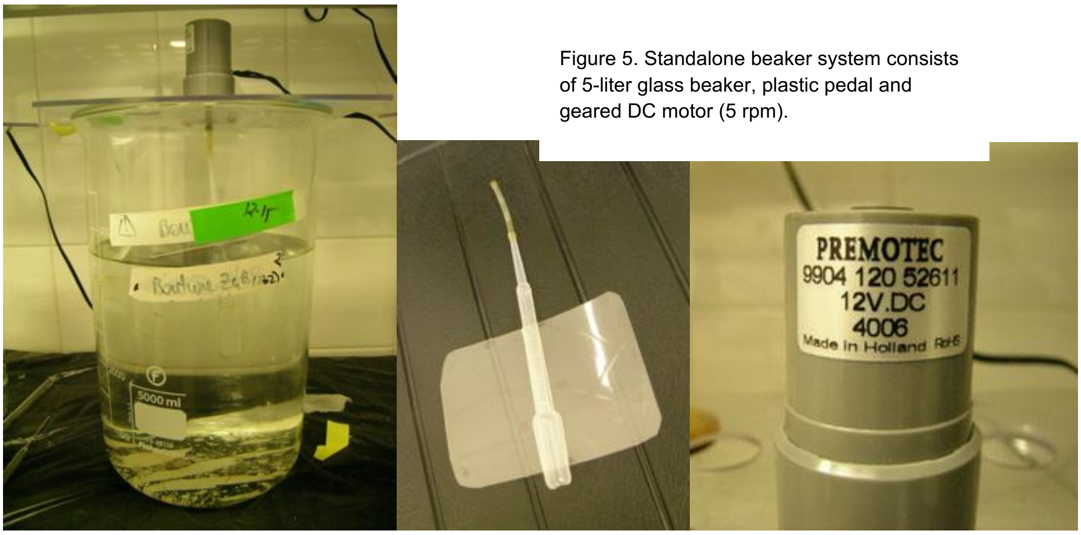


Figure 5. Standalone beaker system consists of 5-liter glass beaker, plastic pedal and geared DC motor (5 rpm).

A large (5 litre) beaker system similar to that described for *Oikopleura* (Bouquet et al., 2009)) can be used for small scale *Clytia* culture for the entire life cycle. It is thus an ideal start-up culture system and also a reliable alternative to the nursery tank. The seawater is stirred horizontally by a plastic or glass paddle (ex. Combining transfer pipette and plastic labware such as a rid of microtiter plate) rotated at 5 rpm by a 12-volt geared electric motor (ex. Premotec 9904 120 52611, 5.0 rpm). Beakers can accommodate any stage of *Clytia*, including polyps and young jellyfish, and are thus suitable for small scale tests. This approach however is more labor-intensive for large scale cultures as the sea water should be replaced manually once or twice a week, and the food concentration must be closely controlled to prevent accumulation of uneaten artemia. Also, medusa growth is significantly slower.

#### Regular maintenance tasks

- Salinity adjustment: every 2~3 days
- Feeding: every day- once or twice.
- Cleaning Kreisel tanks: once every 1~2 weeks, usually when jellyfish are taken out for experiments.
- Cleaning Artemia trap filter in the reservoir: every 2~3 days.
- Sea water change; at least half of the sea water every 2~3 months.
- Cleaning colony tanks and slides: once in 3~4 weeks, usually after collecting juvenile medusa.

### Feeding

We use *Artemia sp*. instar III (corresponding 24 hours after hatching when incubated at between 25 and 30°C) or older (up to 4 days after hatching) nauplii larvae to feed *Clytia.* We feed once a day for polyps or twice a day for medusa, with at least 6 hours between feeding. Unless specified, standard Artemia *sp.* cysts (SepArt, Ocean Nutrition, http://www.oceannutrition.eu/products.aspx?Product=sep-art-artemia-cysts ) were used in this work. Shelf-free Artemia cysts are highly recommended to avoid early medusa death due to bacterial contamination from the cysts. Smaller *Artemia franciscana* (Vinh Chau pond strain, Vietnam) nauplii can be used as an alternative for feeding young medusae and primarily polyps (Van et al., 2014). Detailed instruction is available online (ex. https://www.brineshrimpdirect.com/about-us/articles/hatching-brine-shrimp-cysts/).

#### Preparation of Artemia

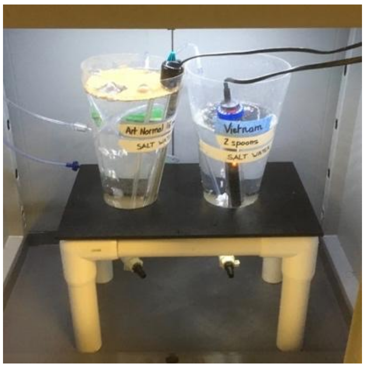


Figure 6. Artemia hatching tank. An incandescent lamp may be used for both lighting and heating purpose. Aeration from the bottom of the tank is critical so that the cysts do not stay in the bottom.

1. Add 1.7 litre of artificial sea water (salinity 30‰) in 2 liter Artemia hatchery cone (ex. https://www.brineshrimpdirect.com/large-hatchery-cone-stand-175-oz-bse ) and install an aquarium heater with thermostat adjusted to a range between 25°C to 30°C. Start aeration and lighting. An incandescent light bulb can be used as a heater.
2. Add up to 16 ml of Artemia cysts on the surface of the seawater.
3. Incubate 24 hours.
4. Stop aeration and heater and wait for 5~15 minutes until newly hatched Artemia larvae sink to the bottom of the cone.
5. Open the valve of the cone and collect Artemia nauplii into a beaker
6. In case of SepArt Artemia, empty and unhatched egg shells are removed using a magnet (http://www.oceannutrition.eu/products.aspx?Product=sep-art-separator ) following the procedure provided by the supplier.
7. Transfer *Artemia* larvae to a sieve (180 µm mesh size) and wash under the running tap water or distilled water for 10~20 seconds. Then rinse with artificial sea water.
8. Incubate *Artemia* larvae in sea water (1~2 liter) at 18~22°C with mild aeration.
9. Start to use them from the following day instar 3 (stage L3) or later (Copf et al., 2003). The color of artemia and swimming pattern can be used to estimate the stage.

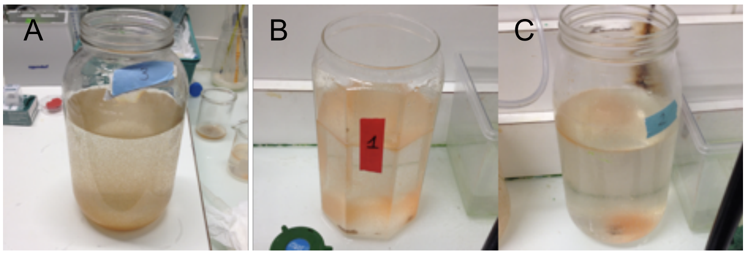


Figure 7 (A) Newly hatched artemia (24 hours of incubation) are yellow/brown color and mostly stay on the bottom. They are not suitable for feeding. (B) Artemia 1 day after hatching (see also supplementary figure) with orange color. They swim actively and some stays near the surface. (C) Artemia 2 days after hatching. Orange becomes less vivid as they develop. The larvae at the stage (B) and (C) actively swim, making a swarm on the bottom or some staying on the surface. Both (B) and (C) are suitable for feeding.

#### Feeding of polyp colonies and medusae

1. Just prior the feeding, wash *Artemia* nauplii in the sieve under running tap water (5~10 seconds) and then resuspend in sea water
2. Add Artemia to the jellyfish/polyp tank using a pipette. Excess feeding is tolerated for Kreisel tanks and polyp tanks. However, the amount of feeing needs to be carefully adjusted depending on the number of mouths to feed for individual tanks (nursery tanks or crystalizing dishes).
3. Check the feeding 30~60 minutes after adding *Artemia* nauplii to the tank and adjust the quantity for next feeding.
4. When medusae are grown in crystalizing dishes, sea water should be changed 3~6 hours after feeding

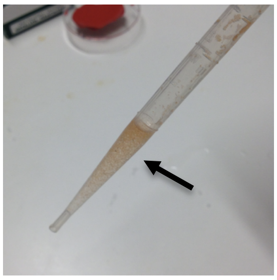

Figure 8. Example of the amount of artemia (arrow) to feed 30 fully grown medusae in a Kreisel tank.

#### Feeding primary polyps with smashed Artemia

"Smashed" *Artemia* nauplii are used for primary polyps, which are significantly smaller than normal polyps.

1. Clean nauplii as described above and transfer a small amount (<1 ml) to an Eppendorf tube
2. Smash nauplii larvae by passing several times through 25 G needle attached to a 1 ml syringe.
3. Transfer smashed nauplii to a small dish and add 5~10 ml sea water and wait for a few minutes until smashed (fragmented) nauplii precipitate on the bottom.
4. Take fragmented nauplii with a Pasteur pipette and slowly apply near the mouth of the primary polyp.

Notes:

- Very young *Artemia* nauplii larvae contain indigestible lipids. With our test, a week of continuous use of instar I larvae killed the colonies by clogging stolon circulation.
- *Artemia* quality has been the most common source of troubles in our aquaculture. Hatched nauplii should thus be systematically washed with tap water and then seawater using a nylon mesh sieve just prior to use.
- Our recent experiences suggested that use of shell-free Artemia cysts reduces early death of medusa due to contamination originating from the egg cysts.

#### Strain duplication by polyp cuttings

*Clytia* strains can be propagated via "cuttings" of the polyp colonies onto fresh slides. If enough polyps are available, we start by putting 5~10 polyps/slide if, as the efficiency is usually low (highly variable between 10% and 50%).

1. Feed the donor polyp colony several hours prior cutting.
2. Cut gastrozooids from the colony at the bottom of the vertical stem (close to the stolon) with microscissors and collect them into a 3 cm dish.
3. Clean the gastrozooids to remove algae by about 10 seconds of vigorous pipetting using a soft plastic pipette.
4. Prepare new glass slides by washing the surface with hot tap water. Label the slide with a diamond pen.
5. Place slides in a petri dish containing sea water to 5~10 mm depth. Transfer gastrozooids onto the slides and leave them undisturbed for one to two nights.
6. Carefully transfer the glass slides into a polyp tank. Feed immediately.

Notes:

- Feeding prior to the cutting favors rapid stolon regrowth.
- We used to scratch the glass surface with sandpaper to help firm attachment of the gastrozooids. We no longer do this because the scratched glass surface also favors algal attachment to the slide and makes cleaning difficult.
- Feeding is critical to keep the transplanted gastrozooids alive. Otherwise the stolon will be extended at the expense of gastrozooids.

#### Colony cleaning

It will be necessary to clean glass plates where the *Clytia* colonies are growing. Red algae will grow in several weeks, depending the intensity and length of aquarium lights. Particles of dead *Artemia* may also precipitate and are accumulated on the slides. These obstacles prevent stolon growth on the glass surface. The surface of glass slides needs to be regularly cleaned so that stolon can be extended, which is critical in long term “colony survival” because there is continuous turnover of polyps and individual polyps (gonozooids) are not immortal. The strategy of cleaning depends on the speed of colony growth and cleaning frequency (Fig.9)

- Conservative cleaning: Clean algae-covered glass surface using wood toothpicks under a stereomicroscope. Try not to touch stolons to keep them as much as possible. Remove dead part of the colony (no cells inside the transparent hydrotheca). This strategy takes long (up to 1 hour for a 75 mm x 50 mm glass slide) but recommended for mutant strains with very slow growth.
- Global cleaning: simply remove the dirtiest parts of the colonies in a glass plate with a fingertip size sponge or a piece of filter pad (see page 3). One can do cleaning without a stereomicroscope, which is however recommended to use to check at the end of cleaning and to identify the dead colonies, which is also to be removed. Usually it takes less than a few minutes for a slide.

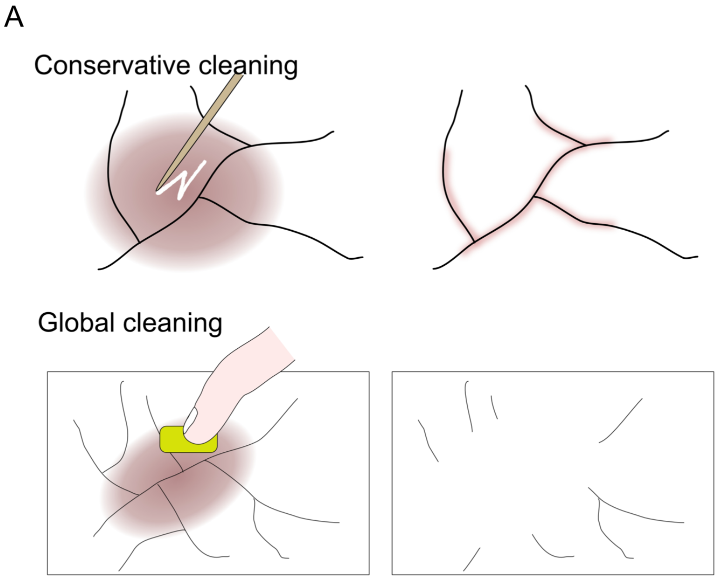


Figure 9. (A) Cleaning strategies. Solid lines represent stolons. Brown paint represent algae growth. See the text above for the detail. (B) Stolon in healthy state. (C) Empty hydrotheca with no stolon cell inside it. This part should be removed together with red algae growing around it. (D) Actively growing stolon tip, which is characterized by "growth cone" morphology. (E) An example of non-growing stolon tip. Stolon is remaining but became very thin. Removing such arrested tip often recovers the growth. Bars= 500 µm

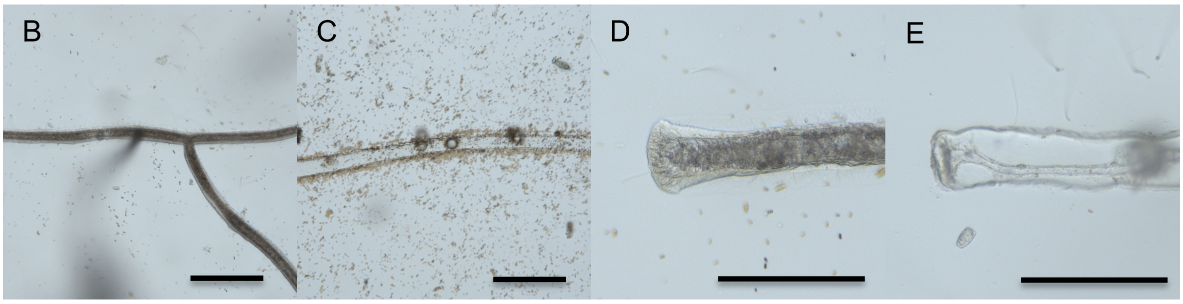


In either case, cleaning helps to stimulate new stolon growth at the cutting edge of the stolon, which may occasionally be arrested the extension with various reasons. It is thus highly recommended to clean the colony frequently (once in a month) before algae spread over the glass surface, even colony can possibly survive more than a year without cleaning.

### In vitro fertilization

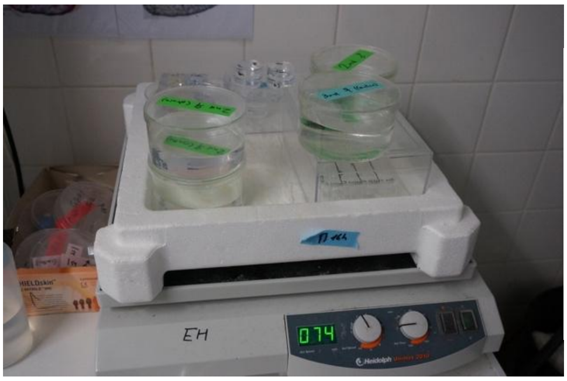

Figure 10. Mature medusae are temporary maintained in crystalizing dishes on a shaker to collect gametes. This can be used to grow out relatively small number (100/dish or less) of juvenile medusae until they became large enough to transfer Kreisel tanks.

Kreisel tanks are maintained under a 24h day-night cycle with a dark period of 3~8 hours. Egg release from the gonad occurs 110- 120 minutes after the light illumination and is highly precise. Sperm release is usually earlier (around 60- 90 minutes after light) and the timing is more variable.  In the protocol below $T_0$ represent the time of dark to light transition (Fig.11).

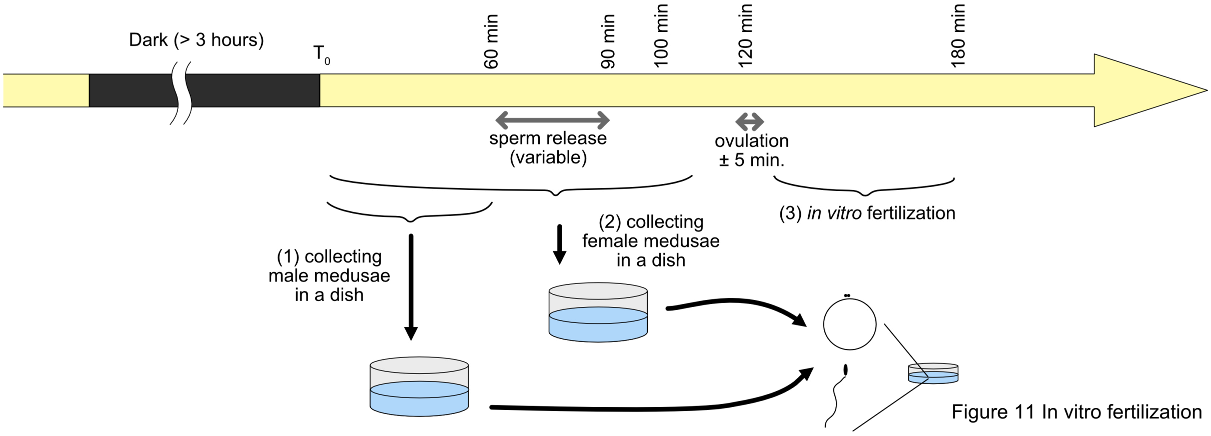


Figure 11 In vitro fertilization

1. At $T_0$+60 min or earlier, transfer male medusae to 10 cm-diameter dishes containing 150~200 ml of seawater (5~20 medusae/dish).
2. At $T_0$+100 min or earlier, transfer female medusae to dishes
   (But not too early to avoid keeping medusa at high density for more than I hour)
3. Once ovulation is complete (at $T_0$+120), transfer medusae back to the Kreisel tank.
4. Eggs can be concentrated in the center of the dish by leaving the dish for 5~10 minutes on the shaker.
5. Eggs are collected in smaller (3.5 cm) dishes for experiments or microinjection
6. Fertilization has to be done by 1 hour after ovulation ($T_0$+180 min). Take sea water from the bottom of the male dish and add to dishes containing eggs. Sperm density should be adjusted following observation under dark-field illumination. Sperm should be visible swimming around the eggs but not present in dense clouds.
7. After fertilization, developing embryos can be selected at the 2- to 8- cell stage (1-2 hours after gamete mixing) or the day after (gastrula stage) and transferred to MFSW and incubated at 20°C. MFSW supplemented with penicillin-streptomycin cocktail (5 unit/ml penicillin and 5

µg/ml streptomycin, Sigma P0781) can be used to prevent bacterial growth that naturally induces uncontrolled metamorphosis. It is sufficient to add penicillin-streptomycin on the day after the fertilization, before planula larvae becomes competent to undergo metamorphosis.

Notes:

- When eggs are fertilized 2 hours (or more) after ovulation, the fertilization rates will be poor and embryonic development will be irregular.
- Mechanical stimulation of male medusae, including pipetting, can break the gonad epidermis causing premature gamete release. It is thus recommended to transfer males into dishes early enough.
- Female jellyfish may be transferred later (90~100 minutes after the light stimulation). This helps to minimize the time spent by medusae in the crystalizing dishes when their number is high (>30/dish).

#### Inducing metamorphosis

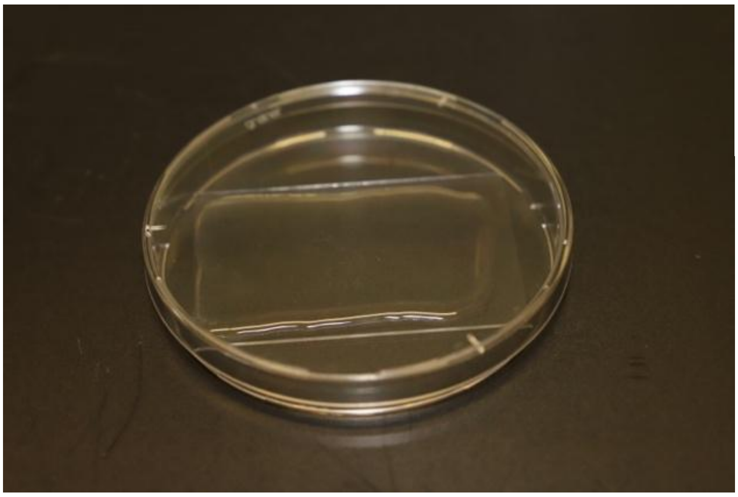

Figure 12. Planula larvae (not visible) being metamorphosized on a large slide (75 mm x 50 mm)

1. Culture planula larvae in MFSW containing penicillin and streptomycin (Sigma P4333, x1/2000 dilution) to avoid bacteria inducing premature metamorphosis.
2. Wash the surface of large glass slides (75 mm x 50 mm) with hot water (40~50°C) or 70% ethanol and wipe off with paper towel. Place them in 100 mm diameter petri dishes.
3. Dilute GLWamide peptide stock solution (1-5 x$10^{-3}$ M in distilled water, stored at -20°C) just prior to use in MFSW (4 ml/large slide). Recommended final concentration for GLWamide6 is 0.5 x $10^{-6}$ M.
4. Using a plastic transfer pipette, carefully pour peptide-containing sea water over the glass slide. Roughly 0.1 ml/$cm^2$ (4 ml for large glass slide).

5. Transfer 2.5~3-day old planulae using a glass pipette. Disperse them as the settlement starts quickly.
6. Cover with a lid to minimize the water evaporation and keep the dish undisturbed.
7. 12~18 hours after peptide treatment, the settled planulae will have formed a “blob” with four-leaf clover shape and be firmly attached to the glass slides. At this stage slides can be transferred to the polyp tanks.
8. Start to feed smashed *Artemia* once the primary polyps have extended tentacles and open mouths. This takes one to several days, timing and efficiency being highly variable.
9. Once the colony contains multiple gastrozooids, it can be fed normally with living *Artemia* nauplii.

#### References


Bouquet, J.-M., Spriet, E., Troedsson, C., Otterå, H., Chourrout, D., and Thompson, E.M. (2009). Culture optimization for the emergent zooplanktonic model organism Oikopleura dioica. J. Plankton Res. *31*, 359–370.

Carré, D., and Carré, C. (2000). Origin of germ cells, sex determination, and sex inversion in medusae of the genus Clytia (Hydrozoa, leptomedusae): the influence of temperature. J. Exp. Zool. *287*, 233–242.

Copf, T., Rabet, N., Celniker, S.E., and Averof, M. (2003). Posterior patterning genes and the identification of a unique body region in the brine shrimp Artemia franciscana. Development *130*, 5915–5927.

Greve, W. (1968). The “Plankton Kreisel” a new device for culturing zooplankton. Marine Biology *1*, 201–203.

Matsakis, S. (1993). Growth of Clytia Spp Hydromedusae (Cnidaria, Thecata) - Effects of Temperature and Food Availability. Journal of Experimental Marine Biology and Ecology *171*, 107–118.

Purcell, J.E., Baxter, E.J., and Fuentes, V.L. (2013). Jellyfish as products and problems of aquaculture. In Advances in Aquaculture Hatchery Technology, (Woodhead Publishing), pp. 404–430.

Van, N.T.H., Van Hoa, N., Bossier, P., Sorgeloos, P., and Van Stappen, G. (2014). Mass selection for small-sized cysts in Artemia franciscana produced in Vinh Chau salt ponds, Vietnam. Aquaculture Research *45*, 1591–1599.
