## Supplemental Figures for "An improved whole life cycle culture protocol for the hydrozoan genetic model *Clytia hemisphaerica*"

*S1. (A and B) Artemia salina after 28 hours of hatching incubation at 30°C, not suitable for feeding. (C) Artemia salina and (D) smaller Artemia franciscana instar 3 nauplius with an additional 24 hours of incubation at room at 20°C, 54 hoursafter start of hatching incubation.(An improved whole life cycle culture protocol for the hydrozoan genetic model Clytia hemisphaerica, Lechable et al.)*

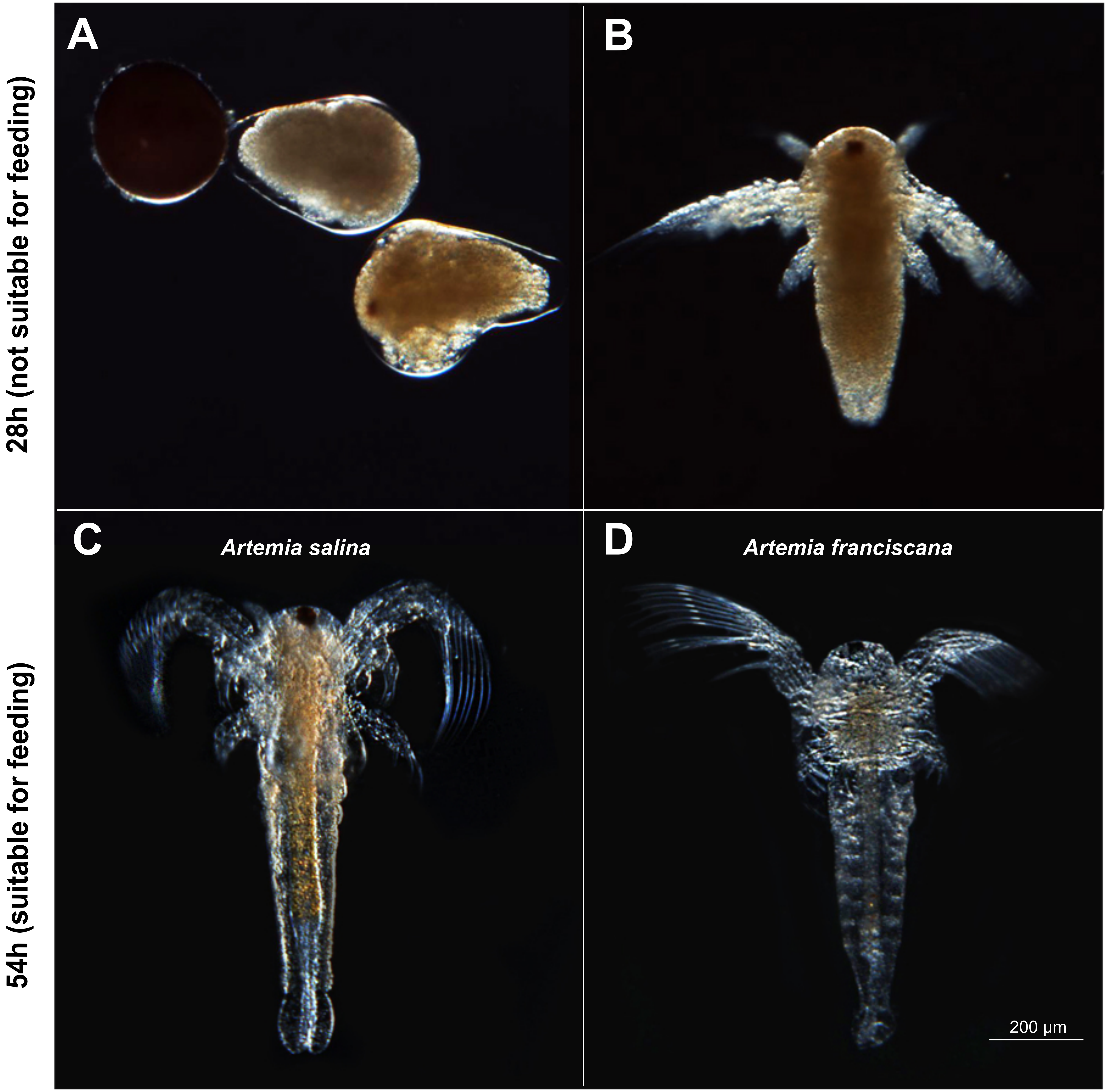

*S2. (A) Proportion of living polyps colonies (N=5 each) after one month incubation at 4°C, 10°C and 18°C without feeding. (B) An example of a polyp after one month at 10°C. (An improved whole life cycle culture protocol for the hydrozoan genetic model Clytia hemisphaerica, Lechable et al.)*

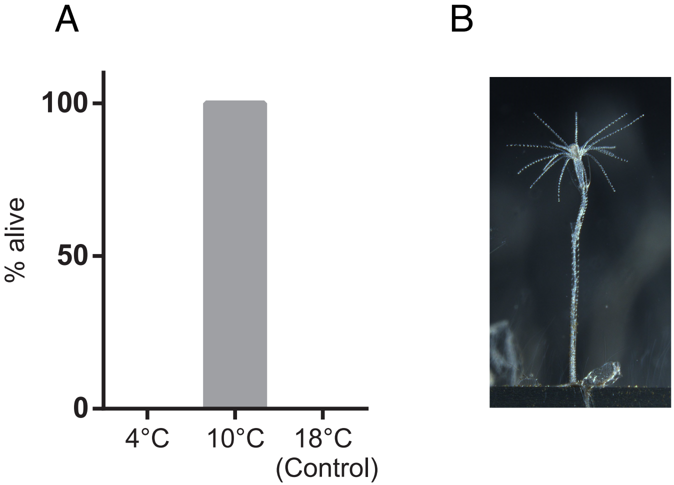

*S3. Disposal of jellyfish and polyp colonies.(An improved whole life cycle culture protocol for the hydrozoan genetic model Clytia hemisphaerica, Lechable et al.)*

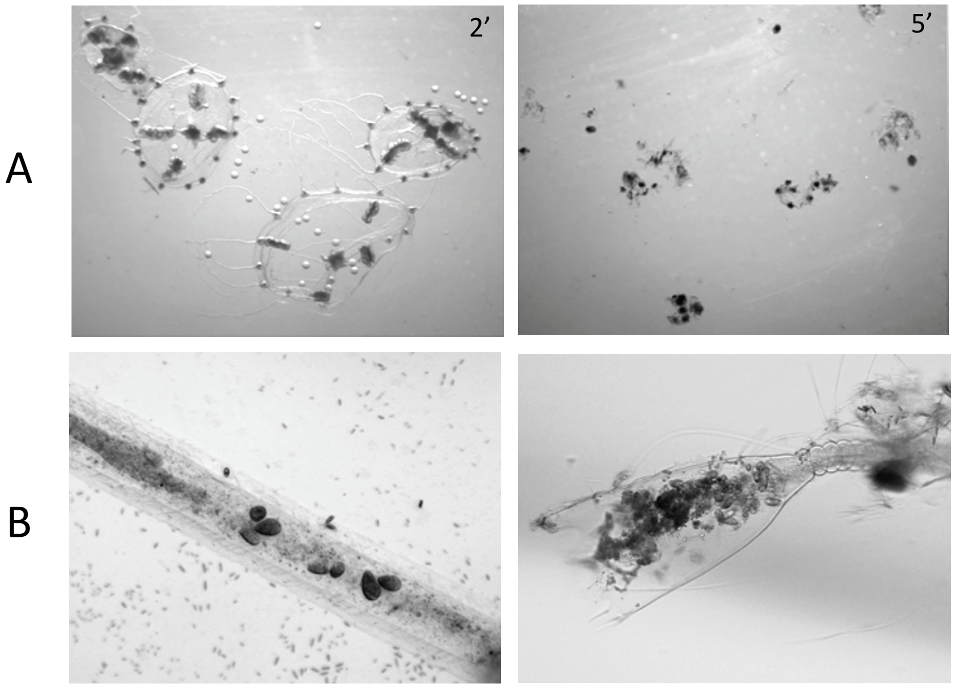

**Supplementary figure**

**S1.** (A and B) *Artemia salina* after 28 hours of hatching incubation at 30°C, not suitable for feeding. Development is not fully synchronized and shows large variation of stages from pre-hatch embryo (A) and instar 1 nauplius (B). Instar 1 nauplius contains high amount of yolk reserve, which is indigestible for *Clytia* polyps and clogs the stolons. (C) *Artemia salina* (body length: 783 +/- 97 μm) and (D) smaller *Artemia franciscana* (body length: 601 +/- 107 μm) instar 3 nauplius with an additional 24 hours of incubation at room at 20°C (total 54 hours after start of hatching incubation). The larval body became transparent and clear orange-colored intestinal tube is visible. This stage of artemia or older are suitable for feeding.

**S2.** (A) Proportion of living polyp colonies (N=5 each) after one month incubation at 4°C, 10°C and 18°C without feeding. (B) An example of a polyp after one month at 10°C.

**S3.** Disposal of jellyfish and polyp colonies. (A) Adult medusae after 2' (left) and 5' (right) and (B) polyp colony by adding 10 volumes of tap water, and disposal of polyp colony after 1h in tap water (S3.B).
